# Upregulation of DNA repair genes and cell extrusion underpin the remarkable radiation resistance of *Trichoplax adhaerens*

**DOI:** 10.1101/2020.12.24.424349

**Authors:** Angelo Fortunato, Alexis Fleming, Athena Aktipis, Carlo C. Maley

**Affiliations:** Arizona Cancer Evolution Center, Arizona State University, 1001 S. McAllister Ave., Tempe, Arizona, USA; Biodesign Center for Biocomputing, Security and Society, Arizona State University, 727 E. Tyler St.,Tempe, Arizona, USA; School of Life Sciences, Arizona State University, 427 East Tyler Mall, Tempe, Arizona, USA; Department of Psychology, Arizona State University, 950 S McAllister Ave, Tempe, Arizona, USA

## Abstract

*Trichoplax adhaerens* is the simplest multicellular animal with tissue differentiation and somatic cell turnover. Like all other multicellular organisms, it should be vulnerable to cancer, yet there have been no reports of cancer in *T. adhaerens*, or any other placozoan. We investigated the cancer resistance of *T. adhaerens*, discovering that they are able to tolerate high levels of radiation damage (218.6 Gy). To investigate how *T. adhaerens* survive levels of radiation that are lethal to other animals, we examined gene expression after the X-ray exposure, finding overexpression of genes involved in DNA repair and apoptosis including the *MDM2* gene. We also discovered that *T. adhaerens* extrudes clusters of inviable cells after X-ray exposure. *T. adhaerens* is a valuable model organism for studying the molecular, genetic and tissue-level mechanisms underlying cancer suppression.

## Introduction

Theoretically, cancer is a disease that can affect all multicellular organisms, and cancer-like phenomena have been observed in all seven branches of the tree of life that independently evolved complex multicellarity[1]. Generally speaking, somatic cells must limit their own proliferation in order for the organism to survive and effectively reproduce. Over the course of 2 billion years, multicellular organisms have evolved many mechanisms to suppress cancer, including control of cell proliferation. Complex multicellularity has evolved independently at least 7 times, and there is evidence of cancer-like phenomena on each of those 7 branches on the tree of life[1]. Although virtually every cell in a multicellular body has the potential to generate a cancer, and that risk accumulates over time, there is generally no association between body size or lifespan and cancer risk, an observation known as Peto’s Paradox[2–5]. This is likely because there has been selective pressure on large, long-lived organisms to evolve better mechanisms to prevent cancer than small, short-lived organisms[6]. This implies that nature has discovered a diversity of cancer suppression mechanisms, which we have only begun to explore for their applications to cancer prevention and treatment in humans[7,8].

We used *Trichoplax adhaerens* (Placozoa) as our model organism for the present study. *T. adhaerens* is the simplest multicellular animal organism ever described (Fig 1A,B). They are also ancient evolutionarily speaking, having diverged from other animals ∼ 800 million years ago[9]. *T. adhaerens* is a disk-shaped, free-living marine organism, 2–3 mm wide and approximately 15 μm high. It is composed of only five somatic cell types, organized into three layers. *T. adhaerens* lack nervous and muscle tissues as well as a digestive system and specialized immune cells. They glide using the cilia of the lower epithelial layer. *T. adhaerens* feed on diatom algae by external digestion. In the laboratory, they reproduce only asexually through fission or budding[10–13] and they feed cooperatively[14]. It is possible to experimentally induce sexual reproduction in the lab, but the embryos do not complete development[10]. It is unknown if *T. adhaerens* reproduce sexually in the wild. *T. adhaerens* can detach from the plate surface when food is depleted and float on the water’s surface. *T. adhaerens* can be collected from the natural world by placing slides in the water column where they are presumably floating[15,16], suggesting that floating is part of the normal behavioral repertoire of *T. adhaerens*.

**Fig 1.**
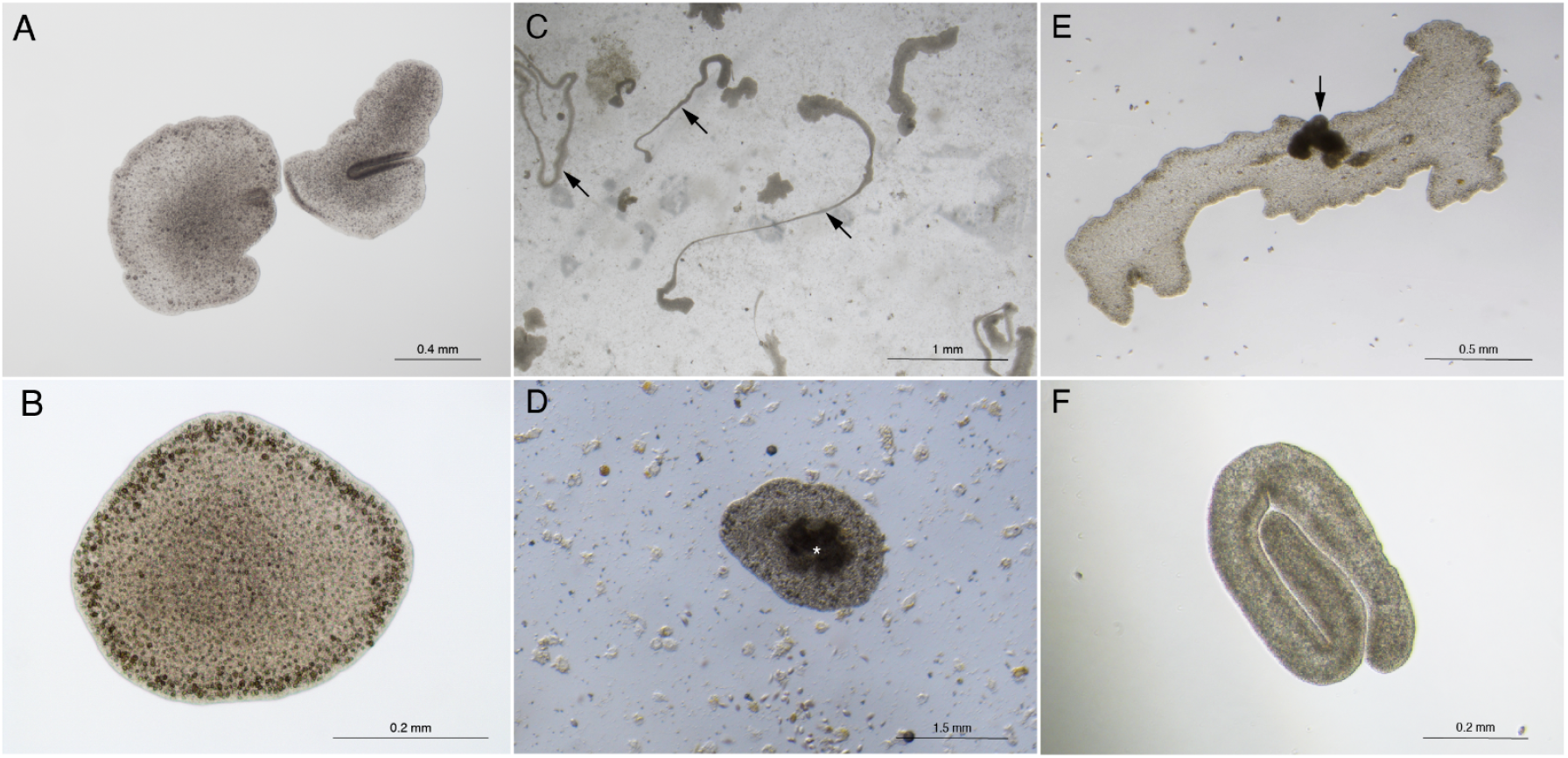
Radiation exposure causes morphological changes in *T. adhaerens*. (A) Untreated specimens of *T. adhaerens*, the animal on the right is folding. (B) Magnification of a single untreated *T. adhaerens*. (C) Sections of the animals can become elongated (arrows), 20 days after 218.6 Gy exposure. (D) Dark tissue mass (asterisk) in the middle of what is either a small animal or extrusion, 70 days after 143.6 Gy exposure. (E) Dark tissue mass projecting from the dorsal epithelium (arrow) of a *T. adhaerens*, 82 days after 143.6 Gy exposure. (F) A folded *T. adhaerens* that is not moving, 36 days after 80 Gy exposure; this animal eventually recovered.

Other invertebrates, such as *Caenorhabditis elegans* and *Drosophila melanogaster* have been useful in molecular biology and the basic sciences[17,18]. However, they are not ideal models for cancer research because they do not have sustained somatic cell turnover, and so do not risk the mutations due to errors in DNA synthesis. In addition, their lifespans are very short, precluding the opportunity to develop cancer. *T. adhaerens*, on the other hand, have somatic cell turnover and very long lifespans - a single organism can reproduce asexually in the lab for decades[19]. Even with these factors that would typically predispose organisms to cancer - cell turnover and long lifespan - there have been no reports of cancer in *T. adhaerens*, despite these organisms having been studied in the laboratory since 1969[20]. In addition, the genome of *T. adhaerens* has been sequenced[21], which enables us to analyze the evolution of cancer genes, detect somatic mutations and quantify gene expression. Despite *T. adhaerens*’ being evolutionarily ancient, most of the known cancer genes in humans have homologs in *T. adhaerens*[21].

It is an open question whether the lack of reports of cancer in *T. adhaerens* is due to a lack of studies or the ability of the animal to resist cancer. We set out to answer this question through exposing *T. adhaerens* to radiation and observing changes in their phenotypes and gene expression. By studying cancer resistance in *T. adhaerens*, it is possible to gain a window into the biological processes and the molecular mechanisms of cancer suppression that likely evolved in the earliest animals.

## Results

We found that *T. adhaerens* are able to tolerate high levels of radiation and are resilient to DNA damage. Exposure to X-rays triggered the extrusion of clusters of cells which subsequently died. We also found that radiation exposure induced the overexpression of genes involved in DNA repair and apoptosis.

### *T. adhaerens* are radiation tolerant

We exposed *T. adhaerens* to different levels of X-ray radiation and counted the number of individuals each day over 8 days after exposure. We then counted them every day for 8 days (S1 Fig). *T. adhaerens* can tolerate 218.6 Gy maximum single dose X-ray exposure. No *T. adhaerens* survived exposure to 256.5 Gy of X-rays. At 218.6 Gy, less than 5% of the *T. adhaerens* survived (measuring the exact percentage is challenging because *T. adhaerens* divide and extrude cells during the experiment but we calculated a lethality of 83.3% after 8 days). These surviving *T. adhaerens* were able to repopulate the culture after 30 days of exposure to 218.6 Gy. We found a statistically significant positive correlation between the doses of radiation (0, 143.6, 181.1, 218.6, 256.5, 294.5 and 332.5 Gy) and the number of *T. adhaerens*, Pearson correlation, r=0.814, p=0.026, calculated as the average of the first 4 days before the beginning of animal death caused by radiation. All the doses, with the exception of 143.6 Gy, cause a sharp decrease in the number of animals by 8 days after the exposure (S1 Fig). We observed morphological and behavioral changes after X-ray exposure, including blisters, changes in the shape of the animals, darker cellular aggregates and extrusion of clusters of cells (Fig 1C-F). These morphological changes were reversible in the animals that survived. *T. adhaerens* that survived also appeared to fully recover.

We found that the total number of discrete *T. adhaerens* entities (including both parents and extruded cells) rapidly increased through budding and fission immediately after X-ray irradiation (Repeated measurement ANOVA, P<0.01, Fig 2A, S2 Fig) and their size significantly decreased (Repeated measurement ANOVA, P<0.0001, Fig 2B), suggesting that the animals extrude cells or divided without physiological cell proliferation to regenerate their original size. After day 7, the total number of *T. adhaerens* in the treated group began to decrease (Fig 2A).

**Fig 2.**
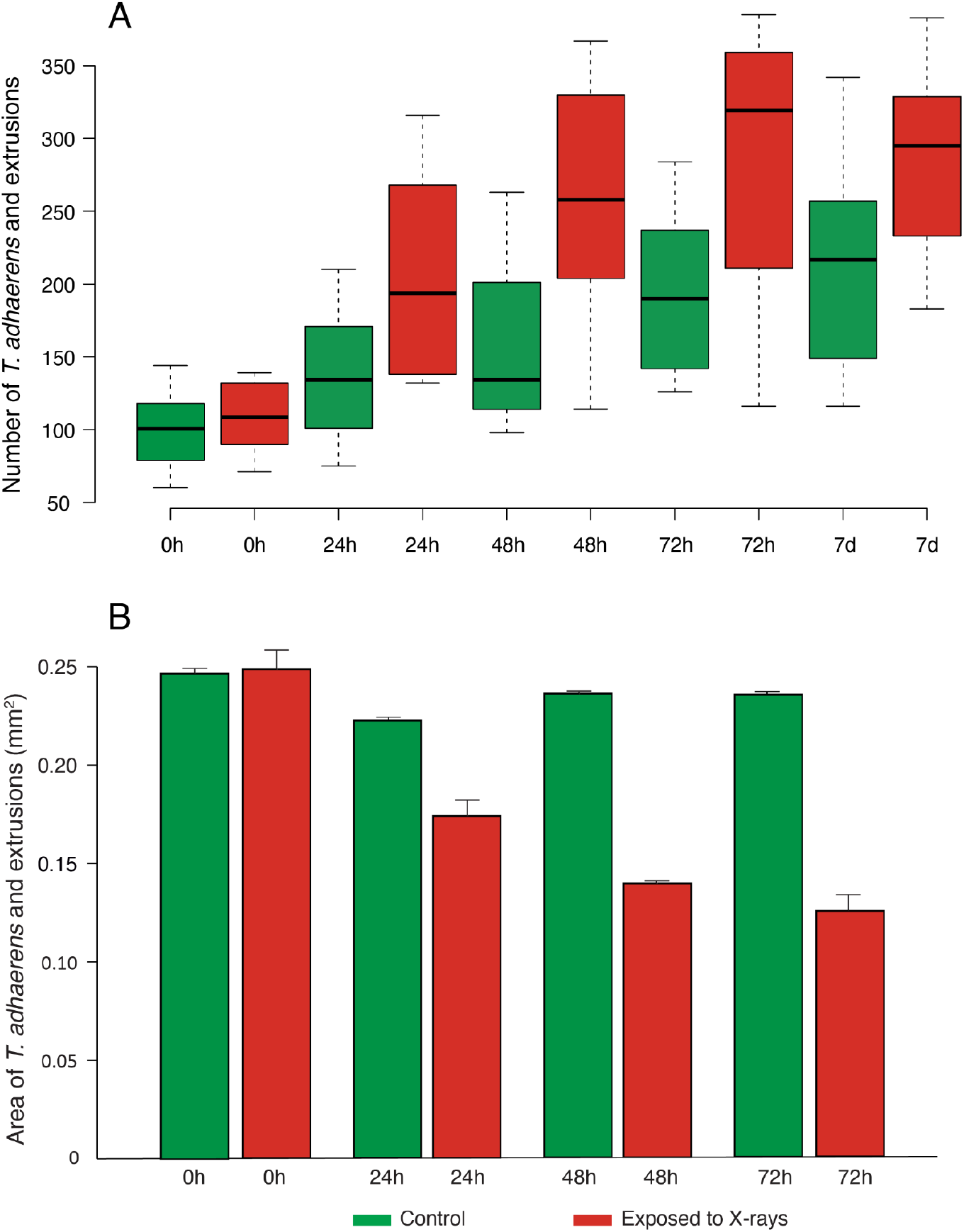
Number and size of X-ray exposed and control *T. adhaerens*. *T. adhaerens* were counted and measured under the microscope and the reported values are a combination of the organisms of all sizes including extrusions. (A) Number of *T. adhaerens* in control (green) and X-ray exposed experimental plates (red) before (0h) exposure, and then 24, 48, 72 hours, and 7 days after exposure to 143.6 Gy of X-rays. Center line =median, box limits indicate the 25th and 75th percentiles. (B) Size of *T. adhaerens* in *T. adhaerens* in control (green) and X-ray exposed experimental plates (red) before (0h), after 24, 48 and 72 hours 143.6 Gy of X-ray exposure. Histograms represent the mean ± s.e.m. (error bars).

### *T. adhaerens* extrude clusters of cells

The extruded bodies (Fig 3) initially are flat and attached to the plate’s bottom but before dying they acquire a spherical shape (S2 Fig). In order to estimate the number of extrusions per animal and to monitor the morphological changes over time, we transferred a single animal per well into 24 well-plates seeded with algae of both control and experimental plates immediately after X-ray exposure. A week after X-ray exposure, the dead extruded buds (65 out of 83 buds) from the experimental animals exceeded the number of dead buds (5 out of 71 buds) in the control (Fisher exact test, P<0.00001). In addition to regular buds, we observed extruded disk-shaped or spherical micro-buds (n=16, ø=182.01μm ± 23.40 s.e.m.) in the experimental plates, but not in the control plates. These micro-buds are only visible at higher magnification, and we did not include them in the number and size measurements of organisms presented in Fig 2.

**Fig 3.**
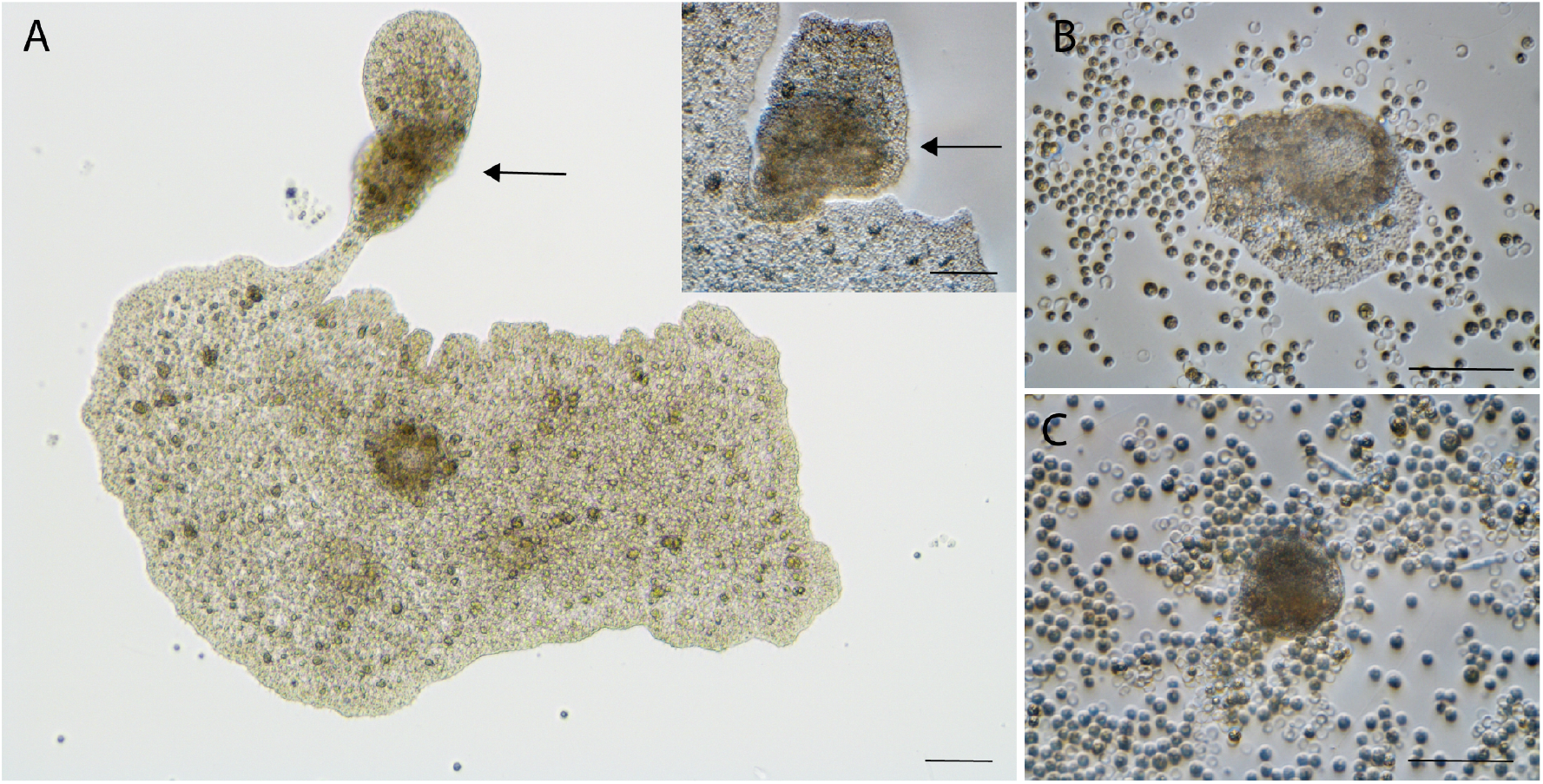
Cell extrusion after radiation exposure. (A) Extrusion (arrows) of brownish putative cancer-like cells; insert, magnification of the same extrusion. (B) The cancer-like cells and the normal cells detached from the main body formed a new animal. The extrusion was observed and isolated 37 days after X-ray exposure. (C) Over time the clear, apparently normal cells of the extruded body reduce in number, leaving only the apparently damaged cells which eventually died. This specimen was exposed to 143.6 Gy X-rays. A: bright field, insert, B and C: differential interference contrast (DIC), scale bars=50μm.

### *T. adhaerens* survive with extensive DNA damage

We tested the animals with a Comet assay and found a catastrophic level of DNA fragmentation soon after a submaximal (143.6 Gy) X-ray exposure (DNA fragmentation: treated= 94.46% ± 0.54 S.D. (n=77 cells), controls=13.83 ± 8.15 S.D. (n=81 cells), Mann-Whitney U Test, P<0.00001, 3 S Fig). The H2AX assay confirmed DNA damage after X-ray exposure (γ-H2AX positive cells: treated= 43.70% ± 13.36 S.D., controls=21.37% ± 4.86 S.D., t-test, P=0.026, Fig. 4S).

**Fig 4.**
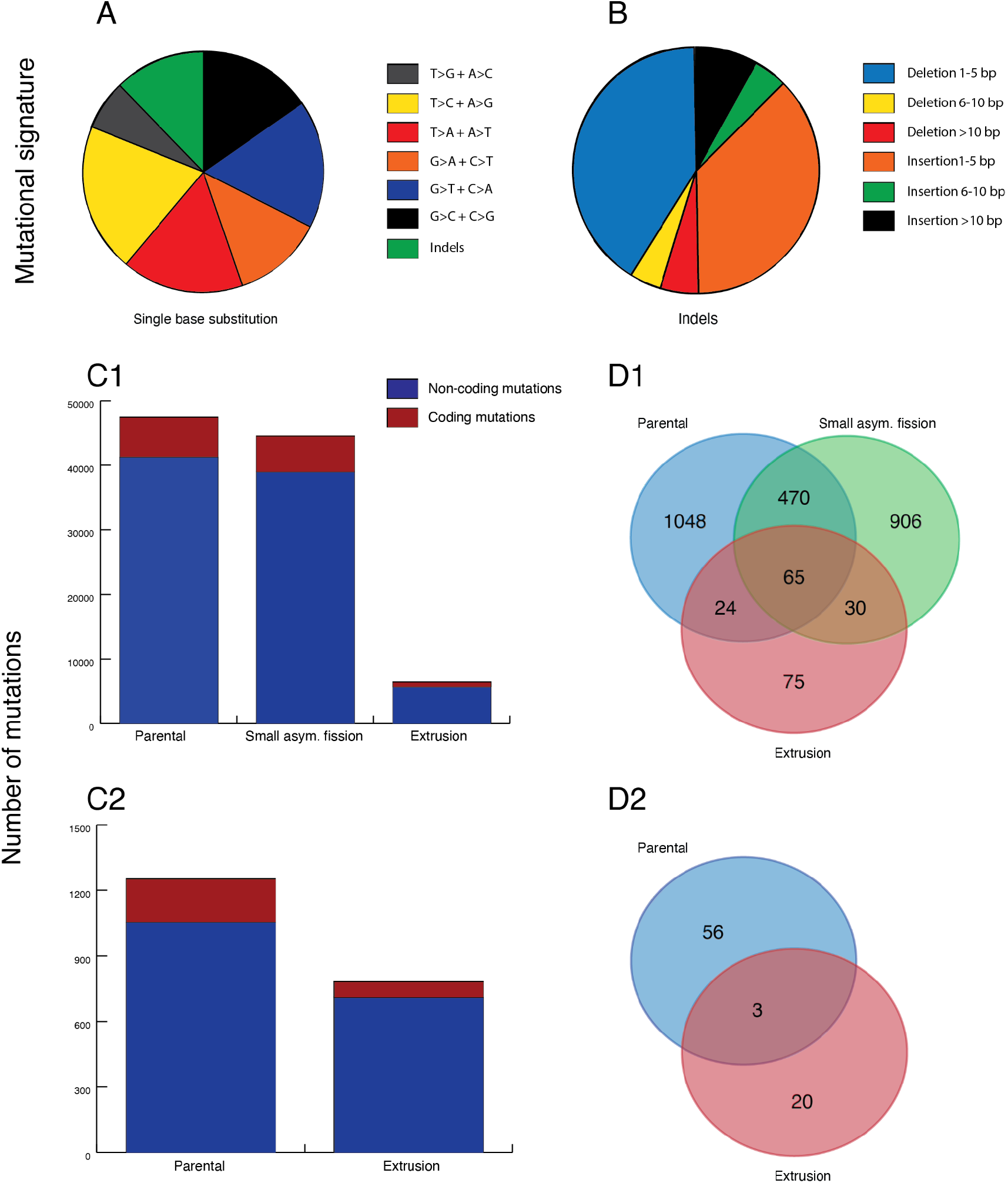
Whole genome sequencing data analysis. (A) Single base substitutions induced by X-ray exposure. (B) Short deletions (40.2%) are statistically more abundant than insertions (36.7%) (paired t-test, p=0.01). (C) Number of mutations in *T. adhaerens* X-ray exposed parental and X-ray exposed extrusion samples. Both types of samples have a high number of mutations. (D) Mutated coding genes overlap between X-ray exposed parental and X-ray exposed extrusion samples, after excluding synonymous mutations. The moderate overlap between X-ray exposed parental and X-ray exposed extrusion samples suggests a different mutational profile between X-ray exposed parental and X-ray exposed extrusion samples. C1 and D1=organism 1, C2 and D2=organism 2, Small asym. fission=Small asymmetric fission.

#### Gene expression changes

We extracted and sequenced RNA from 120 animals 2-hours after the maximal tolerable dose of X-rays (218.6 Gy). We focused on the overexpressed genes because X-ray exposure can generally reduce gene expression. We found 74 genes significantly overexpressed (logFC>2, FDR<0.05) after 2 hours from X-ray exposure (Table 1, S1 Table). Among these, 5 genes with a human ortholog (given in parentheses) are involved in DNA double-strand break repair mechanisms: *TriadG28563* (*RAD52*), *TriadG50031* (*LIG4*), *TriadG53902* (*DCLRE1C*), *TriadG25695* (*RECQL5*), *TriadG61626* (XRCC6). Other genes such as *TriadG55661, TriadG51590, TriadG50243* (*POLB*), *TriadG51591, TriadG28268* (*POLL*), and *TriadG57566* (*LIG3*) are involved in different mechanisms of DNA repair. Interestingly, the *TriadG28470* (*EIF41B*) radioresistant gene[22] is overexpressed after treatment. In addition, we identified up-regulated genes involved in signaling, microtubules activity, transporters, stress response, and other functions. There is marginal or no functional information for 20 of the overexpressed genes (S1 Table).

**Table 1.**
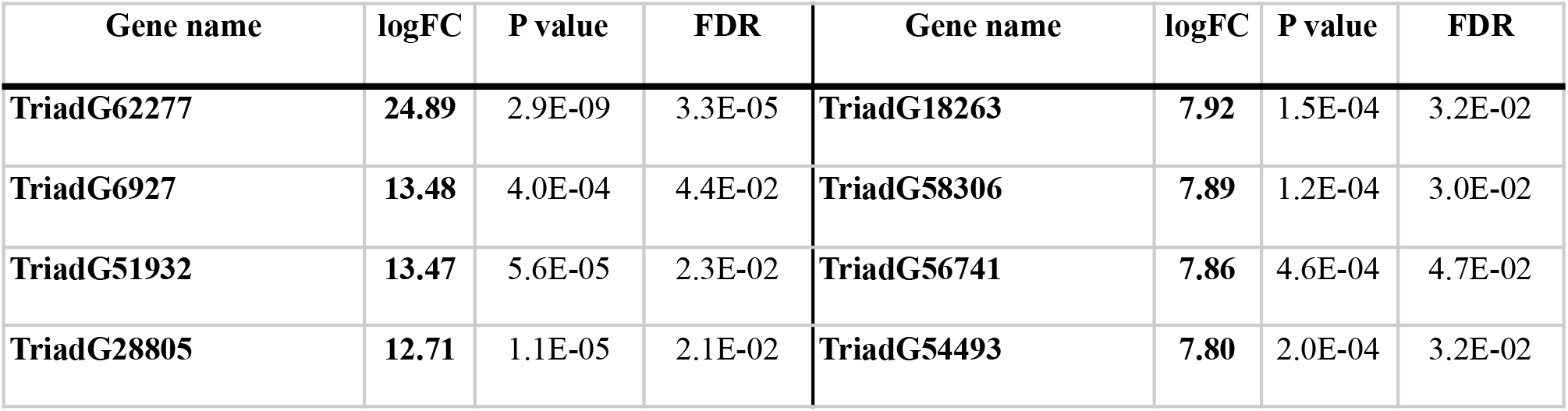
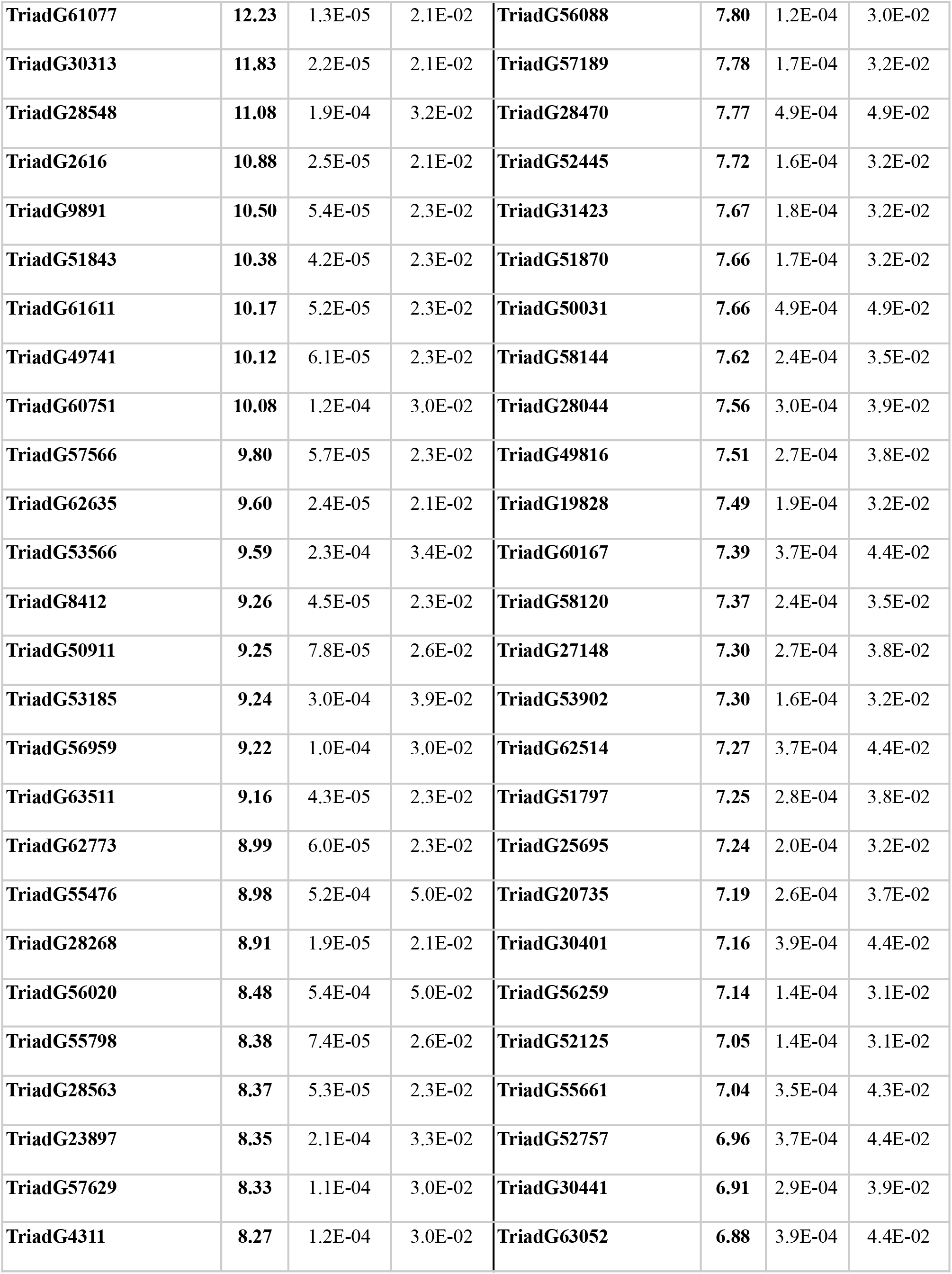
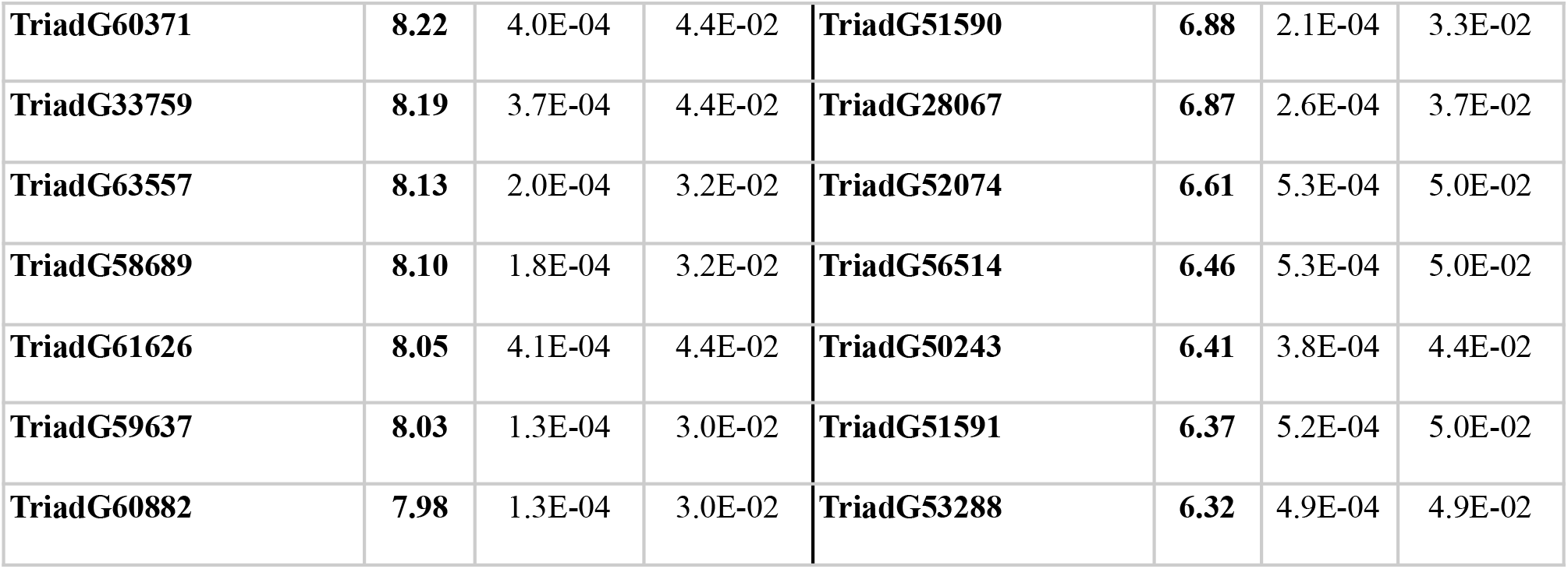
Genes overexpressed after 2 hours following X-ray exposure. Seventeen Five genes are overexpressed in relation to the expression in the control samples, logFC= log_2_ relative fold change. We used the false discovery rate (FDR) correction for multiple comparisons.

| Gene name | logFC | P value | FDR | Gene name | logFC | P value | FDR |
| --- | --- | --- | --- | --- | --- | --- | --- |
| **TriadG62277** | **24.89** | 2.9E-09 | 3.3E-05 | **TriadG18263** | **7.92** | 1.5E-04 | 3.2E-02 |
| **TriadG6927** | **13.48** | 4.0E-04 | 4.4E-02 | **TriadG58306** | **7.89** | 1.2E-04 | 3.0E-02 |
| **TriadG51932** | **13.47** | 5.6E-05 | 2.3E-02 | **TriadG56741** | **7.86** | 4.6E-04 | 4.7E-02 |
| **TriadG28805** | **12.71** | 1.1E-05 | 2.1E-02 | **TriadG54493** | **7.80** | 2.0E-04 | 3.2E-02 |
| **TriadG61077** | **12.23** | 1.3E-05 | 2.1E-02 | **TriadG56088** | **7.80** | 1.2E-04 | 3.0E-02 |
| **TriadG30313** | **11.83** | 2.2E-05 | 2.1E-02 | **TriadG57189** | **7.78** | 1.7E-04 | 3.2E-02 |
| **TriadG28548** | **11.08** | 1.9E-04 | 3.2E-02 | **TriadG28470** | **7.77** | 4.9E-04 | 4.9E-02 |
| **TriadG2616** | **10.88** | 2.5E-05 | 2.1E-02 | **TriadG52445** | **7.72** | 1.6E-04 | 3.2E-02 |
| **TriadG9891** | **10.50** | 5.4E-05 | 2.3E-02 | **TriadG31423** | **7.67** | 1.8E-04 | 3.2E-02 |
| **TriadG51843** | **10.38** | 4.2E-05 | 2.3E-02 | **TriadG51870** | **7.66** | 1.7E-04 | 3.2E-02 |
| **TriadG61611** | **10.17** | 5.2E-05 | 2.3E-02 | **TriadG50031** | **7.66** | 4.9E-04 | 4.9E-02 |
| **TriadG49741** | **10.12** | 6.1E-05 | 2.3E-02 | **TriadG58144** | **7.62** | 2.4E-04 | 3.5E-02 |
| **TriadG60751** | **10.08** | 1.2E-04 | 3.0E-02 | **TriadG28044** | **7.56** | 3.0E-04 | 3.9E-02 |
| **TriadG57566** | **9.80** | 5.7E-05 | 2.3E-02 | **TriadG49816** | **7.51** | 2.7E-04 | 3.8E-02 |
| **TriadG62635** | **9.60** | 2.4E-05 | 2.1E-02 | **TriadG19828** | **7.49** | 1.9E-04 | 3.2E-02 |
| **TriadG53566** | **9.59** | 2.3E-04 | 3.4E-02 | **TriadG60167** | **7.39** | 3.7E-04 | 4.4E-02 |
| **TriadG8412** | **9.26** | 4.5E-05 | 2.3E-02 | **TriadG58120** | **7.37** | 2.4E-04 | 3.5E-02 |
| **TriadG50911** | **9.25** | 7.8E-05 | 2.6E-02 | **TriadG27148** | **7.30** | 2.7E-04 | 3.8E-02 |
| **TriadG53185** | **9.24** | 3.0E-04 | 3.9E-02 | **TriadG53902** | **7.30** | 1.6E-04 | 3.2E-02 |
| **TriadG56959** | **9.22** | 1.0E-04 | 3.0E-02 | **TriadG62514** | **7.27** | 3.7E-04 | 4.4E-02 |
| **TriadG63511** | **9.16** | 4.3E-05 | 2.3E-02 | **TriadG51797** | **7.25** | 2.8E-04 | 3.8E-02 |
| **TriadG62773** | **8.99** | 6.0E-05 | 2.3E-02 | **TriadG25695** | **7.24** | 2.0E-04 | 3.2E-02 |
| **TriadG55476** | **8.98** | 5.2E-04 | 5.0E-02 | **TriadG20735** | **7.19** | 2.6E-04 | 3.7E-02 |
| **TriadG28268** | **8.91** | 1.9E-05 | 2.1E-02 | **TriadG30401** | **7.16** | 3.9E-04 | 4.4E-02 |
| **TriadG56020** | **8.48** | 5.4E-04 | 5.0E-02 | **TriadG56259** | **7.14** | 1.4E-04 | 3.1E-02 |
| **TriadG55798** | **8.38** | 7.4E-05 | 2.6E-02 | **TriadG52125** | **7.05** | 1.4E-04 | 3.1E-02 |
| **TriadG28563** | **8.37** | 5.3E-05 | 2.3E-02 | **TriadG55661** | **7.04** | 3.5E-04 | 4.3E-02 |
| **TriadG23897** | **8.35** | 2.1E-04 | 3.3E-02 | **TriadG52757** | **6.96** | 3.7E-04 | 4.4E-02 |
| **TriadG57629** | **8.33** | 1.1E-04 | 3.0E-02 | **TriadG30441** | **6.91** | 2.9E-04 | 3.9E-02 |
| **TriadG4311** | **8.27** | 1.2E-04 | 3.0E-02 | **TriadG63052** | **6.88** | 3.9E-04 | 4.4E-02 |
| **TriadG60371** | **8.22** | 4.0E-04 | 4.4E-02 | **TriadG51590** | **6.88** | 2.1E-04 | 3.3E-02 |
| **TriadG33759** | **8.19** | 3.7E-04 | 4.4E-02 | **TriadG28067** | **6.87** | 2.6E-04 | 3.7E-02 |
| **TriadG63557** | **8.13** | 2.0E-04 | 3.2E-02 | **TriadG52074** | **6.61** | 5.3E-04 | 5.0E-02 |
| **TriadG58689** | **8.10** | 1.8E-04 | 3.2E-02 | **TriadG56514** | **6.46** | 5.3E-04 | 5.0E-02 |
| **TriadG61626** | **8.05** | 4.1E-04 | 4.4E-02 | **TriadG50243** | **6.41** | 3.8E-04 | 4.4E-02 |
| **TriadG59637** | **8.03** | 1.3E-04 | 3.0E-02 | **TriadG51591** | **6.37** | 5.2E-04 | 5.0E-02 |
| **TriadG60882** | **7.98** | 1.3E-04 | 3.0E-02 | **TriadG53288** | **6.32** | 4.9E-04 | 4.9E-02 |

We focused on two genes: *TP53* (*TriadG64021*) and *MDM2* (*TriadG54791*), the main negative regulator of *TP53*, whose functions in the processes of apoptosis and oncogenesis is well known. *MDM2* and *TP53* genes are well conserved in *T. adhaerens*[23]. RNA-seq analysis suggests that *MDM2* is overexpressed (20-fold), while the expression of *TP53* is similar to its expression in controls. Thus, we conducted additional experiments to investigate *MDM2* and *TP53* genes’ expression, exposing the animals to 218.6 Gy of X-rays. The RNA was extracted at different times after being exposed to X-rays (2, 6, 12, 24, 48 hours). *MDM2* and *TP53* genes’ expression was analyzed by real-time PCR. We found that the expression of *MDM2* was higher (12-fold) after two hours from the beginning of the experiment and decreased over time. On the other hand, the expression of *TP53* was lower and indistinguishable from the controls across all time points (Mann-Whitney test, MDM2 vs control, p<0.05; TP53 vs control, p=NS, Fig. 5S).

#### DNA sequencing of *T. adhaerens* and extruded buds

We sequenced the genomes of two independent pairs of parental X-ray exposed animals and their extruded bodies (as well as a viable, smaller *T. adhaerens* derived from an asymmetric fission of the first parental X-ray exposed animal). We found an average of 1847.8 mutations per Mb (Tab. 2S). In regions of the genome where both X-ray exposed parental samples had at least 10X converge, 0.59% of the detected mutations were shared. Moreover, we found that only 11.9% ± 1.1 S.D. of the total variants detected in all the specimens and 1.9% ± 0.4 S.D. of the coding variants were present in the untreated control population (n=50) of *T. adhaerens*, suggesting that most detected mutations were caused by the X-ray exposure (Fig 4). These percentage values should be understood as maximum overlap between controls and X-ray exposed specimens because not all variants present in the population are necessarily present in the same X-ray exposed specimen. When we described the mutational signature of the treated samples, we found a statistically significant increase of the short deletions in all 5 samples compared to short insertions (paired t-test, p=0.01). This finding is compatible with the mutational profile induced by X-ray exposure (Fig 4). There is a moderate overlap between mutated genes of parental and extrusion samples (Jaccard similarity index, group 1=0.05 (5%), group 2= 0.04 (4%)), suggesting a different mutational profile between parental and extrusion samples. We did not find a statistically significant functional enrichment of mutations in the parental and extrusion samples.

#### DNA sequencing of *T. adhaerens* 2 years after X-ray exposure

We compared 50 random untreated control animals to 50 random animals exposed to X-rays after 2 years. We detected 3637 total mutations (72.7 mutations per animal on average, of which 5 mutations were coding genes, excluding non-synonymous mutations) in the X-ray exposed animals but not in the controls. The number of mutations is strongly reduced compared to the number of mutations detected after 82 and 72 days from X-ray exposure (Fig 4), suggesting that the animals can remove mutations over time. We found that mutated genes with ADP binding function were overrepresented (PANTHER overrepresentation test, fold enrichment=18.14, FDR< 0.0001). All these genes (*TRIADG62071, TRIADG62073, TRIADG62368, TRIADG62451, TRIADG62462, TRIADG62501, TRIADG62549* and *TRIADG62630*) coding for NB-ARC domain-containing protein (Apoptotic Protease-Activating Factor 1 family, PANTHER). We found 68 genes assigned to this family in *T. adhaerens* but only one in humans (*APAF1*), PATHER).

## Discussion

We found that *T. adhaerens* are particularly resilient to DNA damage, which may explain why there have been no reports of cancer in placozoans. Mice die when exposed to 10 Gy of radiation[24,25]. 3–7 Gy of X-rays induces severe DNA damage in mammalian cells[26] and 6 Gy is almost always fatal for humans[27]. In contrast, cancer cell cultures exposed to a cumulative dose of 60 Gy develop radioresistance[28]. What is fascinating about *T. adhaerens* is that despite extensive DNA damage, they survive and fully recover, and in the process, they extrude apparently damaged clusters of cells that eventually die.

### *T. adhaerens* appear to be highly resistant to radiation

We found that some *T. adhaerens* were able to survive extremely high levels (218.6 Gy) of radiation exposure. There are several possible mechanisms that might underlie this radiation resistance. Tardigrades are radioresistant due to mechanisms that prevent DNA damage in the first place[29,30], which seems to be an adaptation to dehydration[31]. Dehydration is unlikely to have been an issue for sea creatures like *T. adhaerens* and their radiation resistance appears not to be due to preventing the DNA damage. In fact, *T. adhaerens* suffer extensive DNA damage from the X-rays, but rely on mechanisms to repair DNA and maintain tissue homeostasis. Also, it is possible that their asexual reproductive strategy of budding reproduction might allow them to make use of many of the same mechanisms to extrude mutated cells in response to radiation exposure.

*T. adhaerens* reproduce vegetatively by repartitioning cells into new individuals. This could expose the population to the risk of spreading cancer cells. Thomas and colleagues [32] propose that the prevention of transmissible cancer could be a factor in the evolution of sexual reproduction, which, combined with immune surveillance, facilitates the detection of a transmissible cancer. They hypothesize that ancient asexual lineages would have had to evolve alternative, efficient mechanisms to prevent cancer from spreading, and note that other ancient asexual organisms (bdelloid rotifers[33] and oribatid mites[34]) are resistant to ionizing radiation and heavy metals. Our results support their hypothesis. Cell extrusion may be one of those mechanisms predicted by Thomas et al. to protect ancient asexual organisms. Because there is no germline/soma distinction in *T. adhaerens*, sexual reproduction could expose an individual to develop gametes from somatic mutated cells or even cancer cells. They also lack an immune system to detect *de novo* or transmissible cancers. Because the disposable soma theory[35] does not apply to placozoans, there may have been strong selective pressure on them to develop alternate mechanisms of cancer suppression.

#### Expression of DNA repair genes and apoptotic pathways increases after radiation exposure

*T. adhaerens* up-regulate genes involved in DNA repair, apoptosis, signaling, microtubule activity, transporters, stress response and radioresistance (Table 1, S1 Table). In particular, our detection of increased expression of the radioresistance gene *TriadG28470* (*EIF41B*) is a nice (positive control) validation of our experimental approach. Interestingly, *TriadG53566* (*SMARCE1*), a gene associated with chromatin remodeling complexes SWI/SNF, is also overexpressed. SWI/SNF complexes are involved in a variety of biological processes, including DNA repair. There is also evidence that *SMARCE1* has a tumor suppressor function[36]. The other genes that were overexpressed after X-ray exposure, with unknown or poorly-known functions may be related to DNA repair, tissue homeostasis or apoptosis. For instance, *Triad28044* is a homolog of the human gene *EMC2*. The function of *EMC2* is not well known in humans but our results suggest that at least one of its functions may be X-ray damage response.

We also found that, after radiation exposure, *MDM2*, the negative regulator of *TP53*, is overexpressed in *T. adhaerens* but *TP53* expression does not increase. This may be an adaptation to prevent catastrophic levels of *TP53*-induced cell death after X-ray exposure, while the animal activates mechanisms of DNA and tissue repair. A possible interpretation of these results is that *MDM2* represses *TP53* activity soon after X-ray exposure. It is also possible that *MDM2* has additional functions related to DNA repair[37]. Although *MDM2* is well conserved in evolution, neither *Caenorhabditis elegans* nor *Drosophila melanogaster* have *MDM2[23]*, suggesting that *T. adhaerens* may be a particularly good model for studying apoptosis.

### *T. adhaerens* extrudes apparently damaged cells that subsequently die

One striking mechanism we observed for dealing with potentially damaged and mutated cells is extrusion of those cells. With the small number of samples and the large number of mutations, we did not have enough statistical power to identify systematic differences in parental and extruded cells. We did detect an over-abundance of mutations in apoptotic pathways, as well as over-expression of *MDM2*. This may be due to natural selection at the cellular level – cells with those mutations and responses would tend to survive, while cells that lacked those mutations and responses probably died.

In *T. adhaerens*, X-ray exposure triggers cell extrusion but the resulting buds are not a form of asexual reproduction. Initially, it is difficult to distinguish extrusion of inviable cells from asexual budding, and so the number of animals seems to increase soon after X-ray exposure. But, as we followed those buds, we found that they almost always die (Fig 3, S2 Fig). This extrusion may be a tissue or organismal strategy to remove damaged cells from the main animal body. We hypothesize that this is a cancer suppression mechanism, extruding pre-malignant cells before they can threaten the organism. This capacity for extrusion of cells might be responsible for the absence of evidence of cancer in *T. adhaerens*.

While the use of extrusion to prevent cancer may seem only relevant to simple organisms, the majority of human cancers arise in epithelial tissues, where extrusion and shedding of damaged cells could be a strategy for eliminating cancerous growths (such as the tissues of the skin and gut). There are hints that similar processes of extrusion of oncogenic cells may be at work in human cancer resistance[38–41]. Apoptotic cells and over-proliferating cells can trigger extrusion[38–41]. The Sphingosine 1-Phosphate pathway contributes to its regulation and is accomplished through cytoskeleton remodeling[42].

The extrusion process is highly conserved in evolution[38–41]. Extrusion is involved in development, initiating cell differentiation, and epithelial-mesenchymal transitions in different organisms ranging from invertebrates to vertebrates[43]. Bacterial infection stimulates shedding, suggesting that cell extrusion is also a defensive mechanism against pathogens; in fact, bacteria can hijack the extrusion molecular mechanisms to invade underlying tissues[43]. Cell cooperation (signaling) and competition are two important factors in extrusion[43]. Cell competition is a cell-elimination process through which cells can eliminate defective (e.g. growth rate, metabolic capacity) adjacent cells. The aberrant activation of signaling pathways in emergent cancer cells can be recognized by normal cells and triggers the elimination of the defective cells.

Cell competition could have a key role in Placozoa because these simple animals have not evolved a complex tissue organization. The extrusion of damaged cells may be a manifestation of cell-cell policing, a process that involves both cell competition and the regulation of cellular cooperation.

Extrusion of damaged cells is an understudied cancer suppression mechanism. At the moment, this process is only partially understood as it can only be studied *in vivo* in intact organisms. The opportunity to study cell extrusion in a simple animal model like *T. adhaerens* allows us to analyze the molecular mechanisms at the base of this process in detail. More broadly, extrusion may allow tissues to defend themselves against neoplastic cells; however, extrusion might also, in some cases, enable the spread of tumoral cellular aggregates in surrounding tissues and in the bloodstream, facilitating the formation of metastasis in advanced tumors. In fact, the metastatic efficiency of tumor cells increases when cells aggregate in multicellular clusters[44]. In this case, it is possible that what was originally a defense mechanism may be subverted by neoplasms in order to metastasize. Understanding this could potentially lead to interventions to help shed pre-cancer cells, to prevent cancer, or alternatively, even suppress this extrusion process to help prevent metastasis.

#### Both *T. adhaerens* and extruded buds have high levels of mutation

The extremely high levels of DNA fragmentation and mutations caused by X-ray radiation suggests that *T. adhaerens* is either very good at repairing DNA or is simply able to tolerate high rates of mutations. The genome sequencing of animals after 2 years from X-ray exposure showed the ability of *T. adhaerens* to survive, apparently without morphological or behavioral changes, harboring 72.7 total mutations in average. The number of mutations is strongly reduced compared to the number of mutations detected after 82 and 72 days from X-ray exposure, suggesting that the animals can remove mutations over time using a combination of cell extrusion and DNA repair mechanisms, suggesting a negative selection of mutated cells and a progressive process of DNA repair. To survive the initial extensive DNA damage, the organisms could activate mechanisms of DNA damage tolerance[45] and repair their DNA through subsequent DNA repair cycles.

The activation of genes involved in repairing DNA double strand breaks through mechanisms of both homologous recombination (e.g. *TriadG28563*) and non-homologous end-joining (e.g *TriadG61626*) had a critical role in the extensive DNA damage recovery. We could hypothesize that other unknown genes overexpressed in response to DNA damage are involved in DNA repair as well. The simplicity of maintaining *T. adhaerens* in culture, the availability of their genome sequence and molecular tools[21] will allow a rapid experimental validation of the function of these genes.

We found a statistically significant enrichment of genes coding for NB-ARC domain-containing protein. There is only one human gene, *APAF1*, with the same protein domain but 68 on *T. adhaerens*. These genes could be involved in the regulation of apoptotic process[46]. The impairment of these genes could have a function in the X-ray damage resilience, inhibiting apoptosis, but at the same time the abundance of these genes could have a role in preventing cancer development. Moreover, the activation of anti-apoptotic genes (e.g. *MDM2*) may prevent damaged cells from dying. *T. adhaerens* may avoid a massive loss of cells due to the extensive damage induced by X-rays by repairing or eliminating the damaged cells over the long term. Importantly, these pathways are well known to be impaired in cancer cells, suggesting that *T. adhaerens* could be a good model to study the mechanisms of carcinogenesis and cancer radioresistance. The low number of samples sequenced do not allow us to draw conclusions pertaining to differences between the parental X-ray exposed specimens and extrusions from the same individuals.

#### Future work and alternative explanations for cell extrusion after radiation exposure

We have suggested that cell extrusion is likely a cancer resistance mechanism in *T. adhaerens*, however, it is possible that extrusion of cells has nothing to do with cancer suppression. One alternative hypothesis that could be tested in future work is that cell extrusion may be a byproduct of *T. adhaerens* asexual fissioning reproductive biology, or even an adaptation to separate into fragments in response to stressors. The reduction of animal size could reduce metabolic demand, mitigating physicochemical stressors[47] and allowing the organism to use more energy to repair cellular damage. Future work could explore these possibilities by deeper sequencing of the parental and extrusion DNA as well as mRNA. RNA expression should reveal if the extrusions are dying cells, neoplastic cells, or (damaged) juvenile organisms, which could be compared to *T. adhaerens* at different stages of reproduction and response to physicochemical stressors. Single-cell DNA sequencing could also reveal if the extrusions are a clonal or a heterogeneous collection of damaged cells.

## Conclusion

Our experiments show that *T. adhaerens* is highly radiation resistant and that radiation exposure causes changes in the expression of genes associated with DNA repair and apoptosis. As a model system, it can potentially be used to identify new genes involved in fundamental processes associated with DNA repair, apoptosis regulation and tissue level cancer protection *in vivo*. Further, *T. adhaerens* is capable of extruding non-viable cells after radiation exposure, suggesting that the process of extrusion might be an important and understudied mechanism of cancer suppression. Together these results show promise for *T. adhaerens* as a model system for studying resilience to radiation exposure as well as the genetic and molecular mechanisms underlying DNA repair and apoptosis.

## Material and methods

### Lab cultures

We grew *T. adhaerens*[48] in glass Petri dishes 100 mm diameter x 20 mm high in 30 ml of artificial seawater (ASW) made in the laboratory by adding 32.5 grams of Instant Ocean sea salt (Prod. n. 77 SS15-10) per liter of distilled water, pH=8, at constant and controlled temperature (23°C) and humidity (60%) with a photoperiod of 14 hrs/10 hrs light/dark cycle in an environmental chamber (Thermo Fisher Scientific, mod. 3940). We fed *T. adhaerens* with diatom algae (*Pyrenomonas helgolandii*) *ab libitum*. Each plate can contain hundreds of animals. When their numbers increase and when food is depleted, *T. adhaerens* detach from the plate surface and float on the water’s surface. We gently collected the floating *T. adhaerens* with a loop and transferred them to new plates, along with 3 ml of ASW from the old plate. This step is required for the animals to successfully grow in the new plates. The animal and algae cultures are assembled in a sterile environment using a biological hood and using sterile materials so that the cultures are protected from parasites and other microorganisms that might interfere with the maintenance of healthy cultures and experiments.

### X-ray exposure

We transferred 50 floating animals in fresh plates with algae 3 days before exposure. On the day of exposure, any floating animals were removed from the plates. We exposed the animals to 160 Gy or 240 Gy using a Rad Source Technologies irradiator (mod. RS-2000 Biological System). Considering X-ray absorbance of the borosilicate glass (Pyrex) plate lid (2 mm) and water in the column above the animals (3 mm), we calculated the actual X-ray exposure of specimens to be 143.6 Gy and 218.6 Gy respectively. To estimate the number of extrusions per animal and to monitor the morphological changes overtime, we transferred a single animal per well into 24 well-plates seeded 7 days before with algae (*P. helgolandii*) of both control and experimental plates. Dose-finding tests suggested that 218.6 Gy is the maximum single dose tolerance for *T. adhaerens*. After 30 days only a few animals (<5%) survived but they repopulated the culture.

### Morphological and morphometric analysis

We observed the animals under a Nikon SMZ1270 dissecting microscope and a Nikon Eclipse Ti-U inverted microscope. We recorded images and videos by using a Nikon DS-Fi2 camera. We counted the treated (n=1085) and control (n=992) *T. adhaerens* and measured the size of the treated (n=1085) and control (n=992) animals present on the 10 plates of control and 10 plates treated replicas using ImageJ software[49]. Images used for morphometric analysis were taken at 20X magnification.

### DNA damage evaluation

We used the silver-stained Comet alkaline assay (Travigen, Cat#4251-050-K)[50,51] to measure the level of DNA fragmentation according to the manufacturer’s specifications and we used ImageJ software[49] to quantify the DNA fragmentation.

### Flow cytometry analysis

The human histone H2AX protein is well conserved in *T. adhaerens* (*TriadG64252*, 82% identity). In particular, serine 139 is present both in human and *T. adhaerens* protein (BLAST[52]). Thus, we used the H2A.X Phosphorylation Assay Kit (Flow cytometry, Millipore, Catalog # 17-344) to detect the level of phosphorylated (serine 139) histone H2AX. We collected 100 animals immediately after the X-ray exposure for each control and experiment replica (3 biological replicas of the experiment) in ASW. We then removed the ASW and we dissociated the cells in cold Mg^++^ and Ca^++^ free PBS with 20 mM EDTA by gently pipetting the collected animals soon after adding the PBS-EDTA buffer. The samples were then kept on ice for 5 minutes to ensure complete dissociation of cells. We processed the dissociated cells according to the manufacturer’s specifications and we quantified the level of phosphorylated histone H2AX using an Attune NxT Flow Cytometer (Invitrogen).

### Gene expression analysis: RNA-seq

We performed the transcriptome analysis using a maximal dose that *T. adhaerens* can tolerate to fully activate the mechanisms of DNA repair and we focused on the early events of DNA damage response. We treated the *T. adhaerens* with 181.1 Gy of X-rays. After 2 hours we extracted the total RNA (RNeasy mini kit, Qiagen, cat. n. 74104) from 40 animals for each of the 3 experimental replicates (total, n=120) and respective controls (total, n=120). After verifying the purity and integrity (RIN= controls: 9.3± 0.2 S.D., experimental 9.2± 0.1 S.D.) of the RNA using an Agilent 2200 TapeStation, part of the extracted RNA was utilized for RNA-seq analysis in order to study the change in genetic expression between controls and experimental specimens at the level of the entire transcriptome. We sequenced the samples using an Illumina NextSeq 500 instrument. We checked the quality of the RNA-seq reads for each sample using FastQC v0.10.1 and we aligned the reads with the reference genome using STAR v2.5.1b (22.68 million reads uniquely mapped on average per sample). Cufflinks v2.2.1 was used to report FPKM values (Fragments Per Kilobase of transcripts per Million mapped reads) and read counts. TPM (Transcripts Per Million) was calculated using R software[53]. We performed the differential expression analysis using the EdgeR package from Bioconductor v3.2 in R 3.2.3. For each pairwise comparison, genes with false discovery rate (FDR) <0.05 were considered significant and log2-fold changes of expression between conditions were reported after Bonferroni correction. We analyzed the differentially expressed genes using Ensembl[54], PANTHER[55] and DAVID [56,57] software.

### Gene expression analysis: real-time PCR

We used the same RNA extracted for the RNA-seq analysis to validate the transcriptome analysis and we extracted the RNA (RNeasy mini kit, Qiagen, Cat. No. 74104) at different times after being exposed to X-rays (2, 6, 12, 24, 48 hours) to study the expression of *TriadG64021* (homolog of the human *TP53* gene) and *TriadG54791* (homolog of the human MDM2 gene). We extracted the RNA from 50 animals coming from 4 different plates exposed to 143.6 Gy X-rays and 4 control plates. Each experiment was repeated thrice. We assessed the RNA integrity through an Agilent 2200 TapeStation system. We retrotranscribed 400 ng of RNA of each specimen using the SuperScript Vilo cDNA synthesis kit (Invitrogen) according to the manufacturer’s protocol. We used the SYBR green fluorescent dye (Power SYBR Green, PCR master mix, Applied Biosystems) to monitor DNA synthesis. We reported the relative expression values for each gene as a ratio of the gene expression level to *TriadT64020* (homolog of the human *GAPDH* gene) expression level in the same sample and normalized for the control level of expression[58,59].

We designed the primers using the software Primer3[60,61] (3 S Table).

### Whole genome sequencing (WGS)

We collected 2 independent groups of specimens: Group 1 is composed by an X-ray exposed specimen (parental), a small animal derived from an asymmetric fission of the X-ray exposed parental animal, and an extrusion derived from the X-ray exposed parental specimen. Group 2 is composed by an X-ray exposed specimen (parental) and an extrusion derived from the same animal. We collected the samples after 82 and 72 days, respectively from the X-ray treatment. Moreover, we collected 50 random untreated control animals and 50 random animals after 2 years from X-ray exposure (143.6 Gy). Both control and X-ray-exposed animals were kept in continuous cultures as described above (lab cultures section). These two groups of animals were pooled in two distinct samples (untreated and X-ray exposed specimens) before DNA extraction. We extracted the DNA using the NucleoSpin Tissue XS kit (Takara, cat.n.740901.50) according to the manufacturer’s specifications.

We generated Illumina compatible Genomic DNA libraries on Agilent’s BRAVO NGS liquid handler using Kapa Biosystem’s Hyper plus library preparation kit (KK8514). We enzymatically sheared the DNA to approximately 300bp fragments, end repaired and A-tailed as described in the Kapa protocol. We ligated Illumina-compatible adapters with unique indexes (IDT #00989130v2) on each sample individually. The adapter ligated molecules were cleaned using Kapa pure beads (Kapa Biosciences, KK8002), and amplified with Kapa’s HIFI enzyme (KK2502). Each library was then analyzed for fragment size on an Agilent’s Tapestation, and quantified by qPCR (KAPA Library Quantification Kit, KK4835) on Thermo Fisher Scientific’s Quantstudio 5 before multiplex pooling and sequencing a 2×100 flow cell on the NovaSeq platform (Illumina) at the Collaborative Sequencing Center.

We loaded 1500pM of the library pool with 1% PhiX for error tracking onto a NovaSeq SP flowcell for 101×8×8×101bp reads. Sequencing was performed using the Illumina NovaSeq 6000 SP Reagent Kit (200 cycles; cat#20040326) on an Illumina NovaSeq 6000. We checked the quality of WGS reads for each sample using FastQC v0.10.1[62] and aligned them to the *T. adhaerens* assembly deposited in DDBJ/EMBL/GenBank as accession number ABGP00000000 using Burrows–Wheeler short-read alignment tool, BWA-MEM version 0.7.15[63]. After alignment, we discovered SNPs and indels following the GATK Best Practices workflow of germline short variant discovery[64]. We pre-processed raw mapped reads by adding read groups, indexing, marking duplicates, sorting, and recalibrating base quality scores. We called the variants using HaplotypeCaller[65]. We discarded them according to their quality score (Q score <30) and coverage (<10X). After discarding those regions, we obtained coverage of 10.1% (parental animal 1), 9.3% (small asymmetric fission from parent 1), 2.7% (extrusion 1), 28.7% (parental animal 2), and 7.6% (extrusion 2) (Tab. 2S). We annotated the variants by SnpEff (version 4.3i)[66].

## Acknowledgments

We would like to thank Arathi Kulkarn and Avalon Yi for help with culturing the animals and assistance during the experiments; Joy Blain, Shanshan Yang, Kristina Buss (ASU Genomics Facility) and Jonathan Scirone for DNA and RNA sequencing.

## Author contributions

A. Fortunato, A.A. and C.C.M. designed the study. A. Fortunato designed and performed the experiments, collected and analyzed the data. A. Fleming was an undergraduate student; she contributed to performing the experiments and to collecting data. A. Fortunato, A.A. and C.C.M. wrote the manuscript.

## Supplementary materials

**1S Fig. Percentage of lethality over time after X-ray exposure**. All radiation doses induce an increased number of *T. adhaerens*. There is a statistically significant positive correlation between the doses of radiation and the number of *T. adhaerens* calculated as the average of the first 4 days before the beginning of animal death caused by radiation (Pearson correlation, r=0.814, p=0.026). All the doses with the exception of 143.6 Gy determine a sharp decrease in the number of animals 8 days after the X-ray exposure. The 8 days final time point is because after 8 days the environmental plates’ condition deteriorates (e.g. reduction of algae, weather quality) and the transferring of animals in fresh plates could compromise their integrity, in particular of the radiation treated ones. Histograms represent the mean± s.e.m. (error bars).

**2S Fig. The extrusions before death acquire a spherical shape (arrows)**; magnification 40X.

**3S Fig. Representative images of the comet assay measuring DNA strand breaks**. (A) The controls have few nuclei showing DNA fragmentation. (B) In contrast, animals exposed to 143.6 Gy of X-rays have many more nuclei with extensive fragmentation of their DNA (Mann-Whitney U Test, P<0.0001). The arrows indicate examples of a “comet” with the nucleus containing unfragmented DNA and the electrophoretic migration of fragmented DNA (tail, shown in the inset of panel B).

**4S Fig. H2AX phosphorylation DNA damage assay**. We confirmed DNA damage using the H2AX phosphorylation assay, controls (A, B), experimental (C, D) cells. The solid line shows the region of cell-derived fluorescence signals.

**5S Fig. Change in *TP53* and *MDM2* gene expression in *T. adhaerens* after X-ray exposure**. Each experiment was repeated thrice (Mann-Whitney test, MDM2 vs control, p<0.05; TP53 vs control, p=NS; TP53 vs MDM2, paired t-test, P<0.05). Histograms represent the mean (log_10_ fold) ± s.e.m. (error bars).

**S1 Table. Annotations of genes overexpressed after X-ray exposure**. We used the DAVID software to annotate the overexpressed genes. This table reports the name of the gene, the biological function, the cellular component, the molecular function, protein description (INTERPRO), the pathway of appurtenance (KEGG) and the SMART annotation if available.

**S2 Table. Coverage of whole genome sequencing**. There is a substantial difference between the coverage of the parental and extruded samples. This is mostly caused by the limited-dying number of cells available in the extrusions.

**S3 Table. PCR primer sequences**. Sequences of primers (5’-3’) of *T. adhaerens* genes and their human orthologs (as indicated in parentheses), *TriadT64020* (*GAPDH*), *TriadG64021* (*TP53*) and *TriadG54791* (*MDM2*).

