## Supplementary figures and tables for "Upregulation of DNA repair genes and cell extrusion underpin the remarkable radiation resistance of *Trichoplax adhaerens*"

**Supplementary materials.**

**Figure 1S**

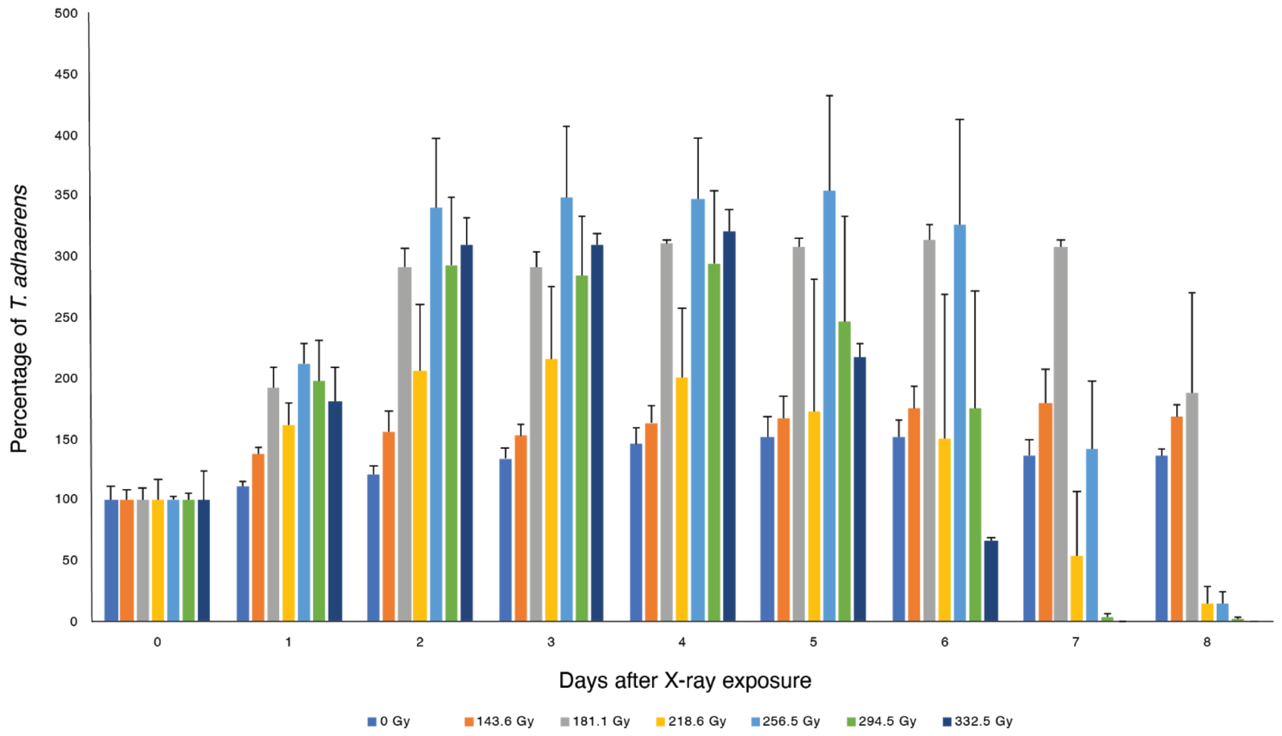


**Figure 1S.** Percentage of lethality over time after X-ray exposure. All radiation doses induce an increased number of *T. adhaerens.* There is a statistically significant positive correlation between the doses of radiation and the number of *T. adhaerens* calculated as the average of the first 4 days before the beginning of animal death caused by radiation (Pearson correlation, r=0.814, p=0.026). All the doses with the exception of 143.6 Gy determine a sharp decrease in the number of animals during the first 8 days after the exposure. The 8 days final time point is because after 8 days the environmental plates' condition deteriorates (e.g. reduction of algae, weather quality)

and the transferring of animals in fresh plates could compromise their integrity, in particular of the radiation treated ones. Histograms represent the mean± s.e.m. (error bars).

**Figure 2S.**

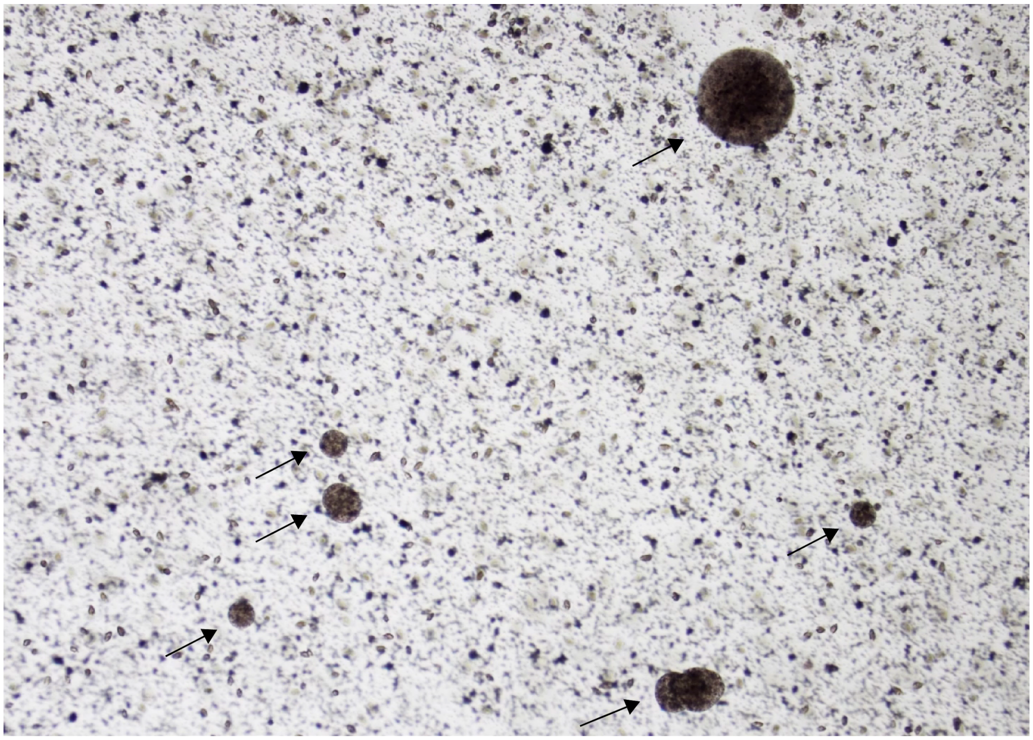

**Figure 2S.** The extrusions before death acquire a spherical shape (arrows); magnification 40X.

**Figure 3S.**

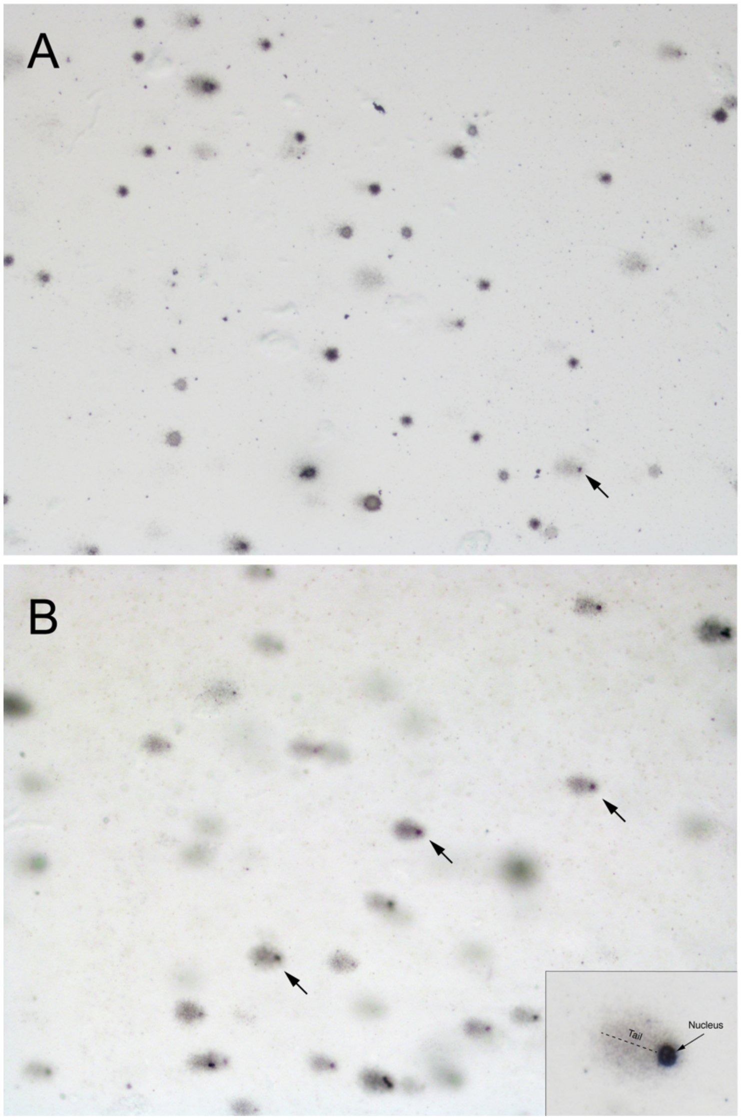


.

**Figure 3S.** Representative images of the comet assay measuring DNA strand breaks. **A.** The controls have few nuclei showing DNA fragmentation. **B.** In contrast, animals exposed to 143.6 Gy of X-rays have many more nuclei with extensive fragmentation of their DNA (Mann-Whitney U Test, P<0.0001). The arrows indicate examples of a "comet" with the nucleus containing unfragmented DNA and the electrophoretic migration of fragmented DNA (tail, shown in the inset of panel B).

**Figure 4S.**

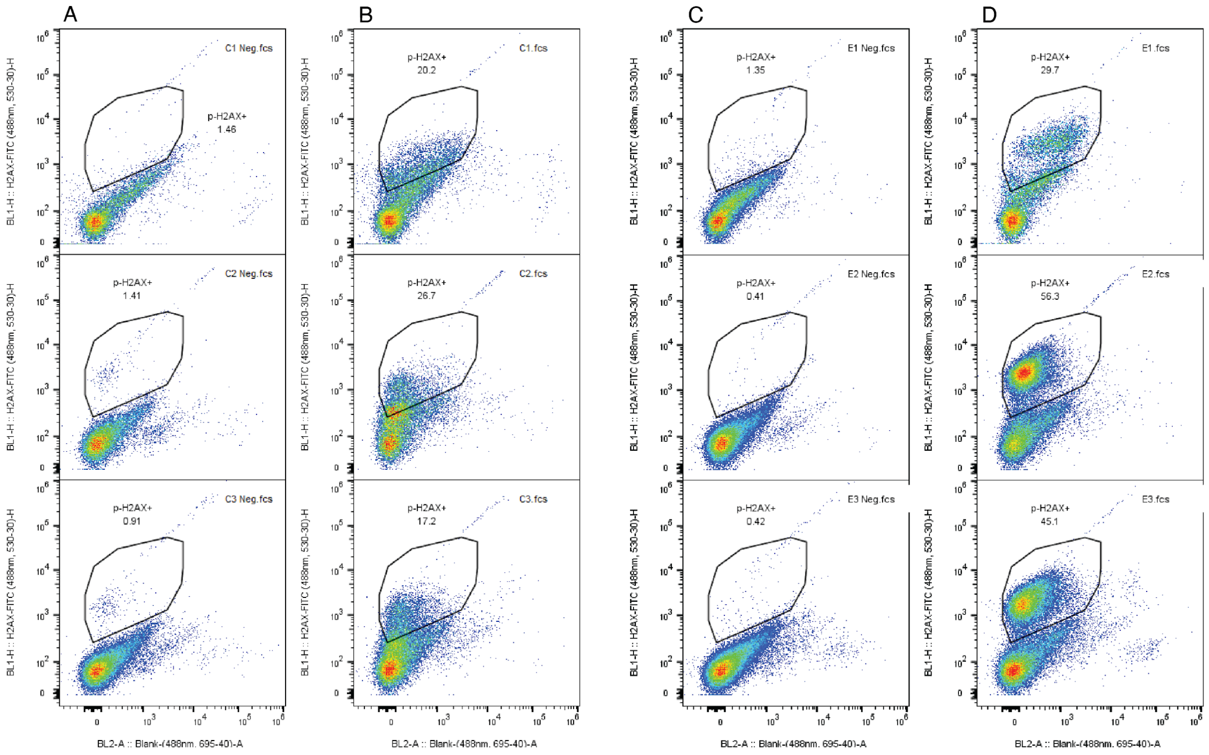


**Figure 4S.** We confirmed DNA damage using the H2AX phosphorylation assay, controls (A, B), experimental (C, D) cells. The solid line shows the region of cell-derived fluorescence signals.

**Figure 5S.**

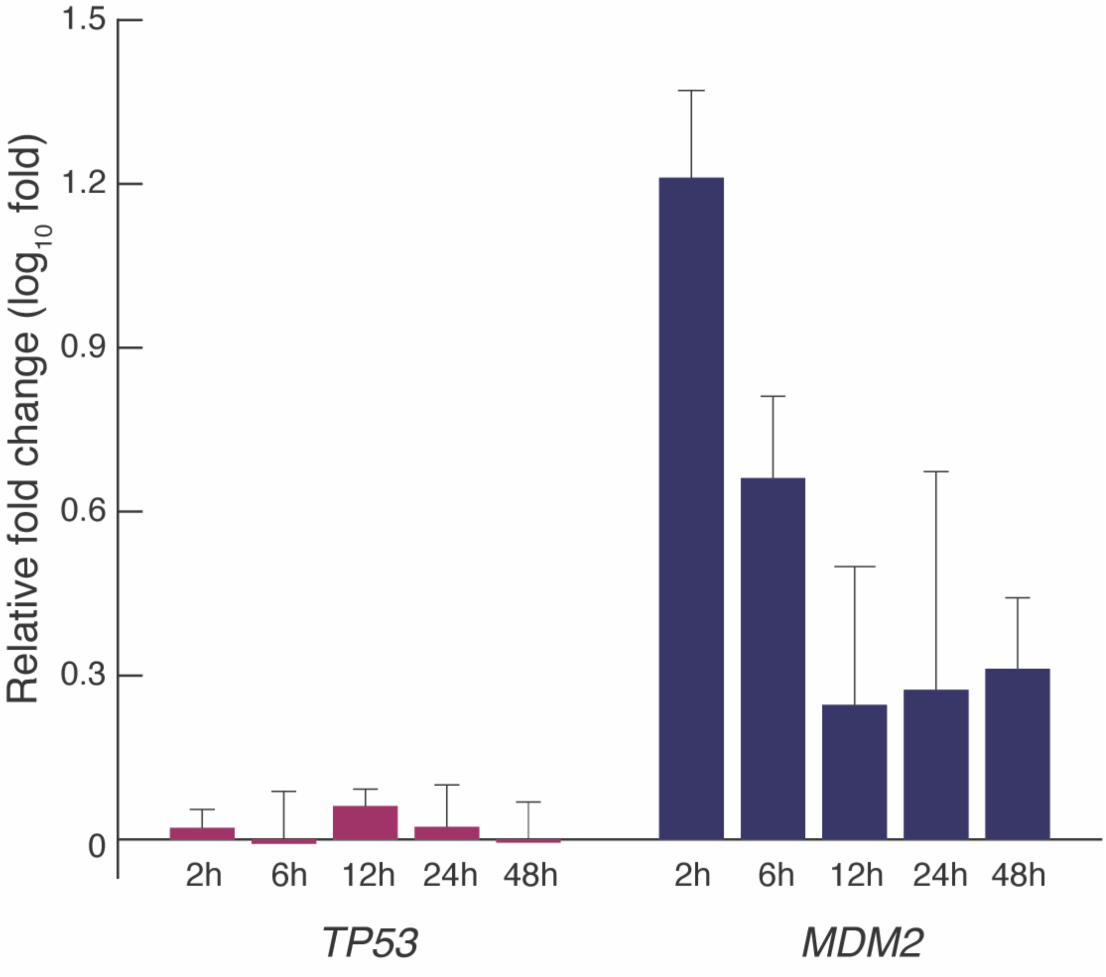
Relative fold change ($log_{10}$ fold)
1.5
1.2
0.9
0.6
0.3
0
2h
6h
12h
24h
48h
TP53
2h
6h
12h
24h
48h
MDM2

**Figure 5S.** Change in *TP53* and *MDM2* gene expression in *T. adhaerens* after X-ray exposure. Each experiment was repeated thrice (Mann-Whitney test, MDM2 vs control, $p<0.05$; TP53 vs control, p=NS; TP53 vs MDM2, paired t-test, $P<0.05$). Histograms represent the mean ($\log_{10}$ fold) ± s.e.m. (error bars).

**Table 1S.** Annotations of genes overexpressed after X-ray exposure. We used the DAVID software to annotate the overexpressed genes. This table reports the name of the gene, the biological function, the cellular component, the molecular function, protein description (INTERPRO), the pathway of appurtenance (KEGG) and the SMART annotation if available.

| Gene & annotation methods | Description |
|---|---|
| **TriadG50911** | |
| GOTERM Biological Process | regulation of transcription from RNA polymerase II promoter, |
| GOTERM Cellular Component | nucleus, |
| GOTERM Molecular Function | transcription factor activity, sequence-specific DNA binding |
| INTERPRO | Homeodomain, Homeodomain-like, Homeobox |
| SMART | HOX, |
| **TriadG63557** | |
| GOTERM Biological Process | regulation of transcription from RNA polymerase II promoter, transcription elongation from RNA polymerase II promoter, protein transport, histone deubiquitination, poly(A)+ mRNA export from nucleus, positive regulation of transcription, DNA-templated, |
| GOTERM Cellular Component | SAGA complex, nuclear pore, transcription export complex 2, DUBm complex, |
| GOTERM Molecular Function | chromatin binding, transcription coactivator activity, |

| | |
|---|---|
| INTERPRO | Transcription factor, enhancer of yellow 2, |
| **TriadG58120** | Uknown |
| **TriadG18263** | |
| GOTERM Biological Process | cell redox homeostasis, |
| GOTERM Cellular Component | cell, |
| GOTERM Molecular Function | oxidoreductase activity, |
| INTERPRO | Thioredoxin-like fold, Redoxin, Thioredoxin domain, |
| KEGG_PATHWAY | Peroxisome, |
| **TriadG19828** | |
| GOTERM Cellular Component | integral component of membrane, |
| INTERPRO | Drug/metabolite transporter, Solute carrier family 35 member F3/F4, |
| **TriadG20735** | |
| INTERPRO | Vacuolar sorting protein 9, |
| SMART | VPS9, |
| **TriadG23897** | |
| GOTERM Biological Process | sterol metabolic process, |
| GOTERM Cellular Component | integral component of membrane, |
| GOTERM Molecular Function | cholestenol delta-isomerase activity, |
| INTERPRO | Emopamil-binding, |
| KEGG_PATHWAY | Steroid biosynthesis, Metabolic pathways, |
| **TriadG25695** | |
| GOTERM Biological Process | double-strand break repair via homologous recombination, |
| GOTERM Cellular Component | nucleus, chromosome, cytoplasm, |
| GOTERM Molecular Function | DNA binding, ATP binding, four-way junction helicase activity, ATP-dependent 3'-5' DNA helicase activity, |

| | |
|---|---|
| INTERPRO | Helicase, C-terminal, DNA helicase, ATP-dependent, RecQ type, DNA/RNA helicase, DEAD/DEAH box type, ATP-binding domain, P-loop containing nucleoside triphosphate hydrolase, |
| SMART | DEXDc, HELICc, |
| **TriadG2616** | |
| GOTERM Cellular Component | kinesin complex, |
| GOTERM Molecular Function | microtubule motor activity, |
| INTERPRO | Kinesin light chain, Tetratricopeptide-like helical, Tetratricopeptide repeat-containing domain, |
| SMART | TPR, |
| **TriadG27148** | |
| GOTERM Cellular Component | integral component of Golgi membrane, integral component of endoplasmic reticulum membrane, |
| GOTERM Molecular Function | 3'-phosphoadenosine 5'-phosphosulfate transmembrane transporter activity, |
| INTERPRO | UAA transporter, |
| **TriadG28044** | |
| INTERPRO | Tetratricopeptide-like helical, Tetratricopeptide repeat-containing domain, Tetratricopeptide repeat, |
| SMART | TPR, |
| **TriadG28067** | |
| GOTERM Cellular Component | integral component of membrane, |
| GOTERM Molecular Function | N-acetyltransferase activity, |
| INTERPRO | GNAT domain, Acyl-CoA N-acyltransferase, |
| **TriadG28268** | |
| GOTERM Biological Process | DNA repair, |
| GOTERM Cellular Component | nucleus, |

| | |
|---|---|
| GOTERM Molecular Function | DNA binding, DNA-directed DNA polymerase activity, |
| INTERPRO | DNA polymerase, family X, beta-like, DNA-directed DNA polymerase X, DNA polymerase beta-like, DNA polymerase lambda, fingers domain, DNA polymerase family X lyase domain, |
| KEGG_PATHWAY | Base excision repair, Non-homologous end-joining, |
| SMART | POLXc, |
| **TriadG28470** | |
| GOTERM Cellular Component | cytoplasm, |
| GOTERM Molecular Function | translation initiation factor activity, |
| INTERPRO | Translation Initiation factor eIF- 4e, Eukaryotic translation initiation factor 4E (eIF-4E), |
| KEGG_PATHWAY | RNA transport, mTOR signaling pathway, |
| **TriadG28548** | |
| INTERPRO | BolA protein, |
| PIR_SUPERFAMILY | Stress-induced morphoprotein, BolA type, |
| **TriadG28563** | |
| GOTERM Biological Process | DNA recombinase assembly, double-strand break repair via single-strand annealing, |
| GOTERM Cellular Component | nucleus, |
| INTERPRO | DNA recombination/repair protein Rad52, Rad52/22 double-strand break repair protein, |
| KEGG_PATHWAY | Homologous recombination, |
| **TriadG28805** | |
| GOTERM Biological Process | ubiquitin-dependent protein catabolic process, mitotic cell cycle checkpoint, |
| GOTERM Cellular Component | nucleus, |
| GOTERM Molecular Function | ubiquitin-protein transferase activity, |
| **TriadG30313** | |
| GOTERM Biological Process | mesoderm development, regulation of lipid storage, |

| | |
|---|---|
| GOTERM Molecular Function | hydrolase activity, |
| INTERPRO | Alpha/beta hydrolase fold-1, Epoxide hydrolase-like, Phosphoglycolate phosphatase, domain 2, HAD-like domain, |
| KEGG_PATHWAY | Arachidonic acid metabolism, Metabolic pathways, Peroxisome, |
| **TriadG30401** | |
| GOTERM Molecular Function | cysteine-type peptidase activity, |
| INTERPRO | Peptidase C48, SUMO/Sentrin/Ubl1, |
| **TriadG30441** | |
| GOTERM Biological Process | cellular copper ion homeostasis, response to oxidative stress, intracellular copper ion transport, |
| GOTERM Cellular Component | cytoplasm, |
| GOTERM Molecular Function | copper chaperone activity, |
| INTERPRO | Heavy metal-associated domain, HMA, |
| **TriadG31423** | |
| GOTERM Cellular Component | integral component of membrane, |
| INTERPRO | Sec20, |
| KEGG_PATHWAY | SNARE interactions in vesicular transport, |
| **TriadG33759** | |
| GOTERM Cellular Component | integral component of membrane, |
| GOTERM Molecular Function | ATP binding, |
| INTERPRO | Chaperonin ClpA/B, Torsin, P-loop containing nucleoside triphosphate hydrolase, |
| **TriadG4311** | |
| GOTERM Biological Process | cell surface receptor signaling pathway, G-protein coupled receptor signaling pathway, |
| GOTERM Cellular Component | integral component of plasma membrane, |
| GOTERM Molecular Function | G-protein coupled receptor activity, |
| INTERPRO | G protein-coupled receptor, rhodopsin-like, GPCR, rhodopsin-like, 7TM, |

| | |
|---|---|
| **TriadG49741** | |
| INTERPRO | WD40 repeat, WD40/YVTN repeat-like-containing domain, WD40-repeat-containing domain, G-protein beta WD-40 repeat, |
| KEGG_PATHWAY | mRNA surveillance pathway, |
| SMART | WD40, |
| **TriadG49816** | |
| INTERPRO | Tetratricopeptide-like helical, Tetratricopeptide repeat-containing domain, Tetratricopeptide repeat, |
| SMART | TPR, |
| **TriadG50031** | |
| GOTERM Biological Process | DNA replication, nucleotide-excision repair, DNA gap filling, double-strand break repair via nonhomologous end joining, DNA recombination, DNA ligation involved in DNA repair, |
| GOTERM Cellular Component | cytoplasm, DNA-dependent protein kinase-DNA ligase 4 complex, DNA ligase IV complex, |
| GOTERM Molecular Function | DNA binding, DNA ligase (ATP) activity, ATP binding, |
| INTERPRO | DNA ligase, ATP-dependent, BRCT domain, DNA ligase, ATP-dependent, N-terminal, DNA ligase, ATP-dependent, C-terminal, DNA ligase, ATP-dependent,, OB-fold, DNA ligase, ATP-dependent, DNA ligase IV, |
| KEGG_PATHWAY | Non-homologous end-joining, |
| SMART | BRCT, |
| **TriadG50243** | |
| GOTERM Biological Process | DNA repair, |
| GOTERM Cellular Component | nucleus, |
| GOTERM Molecular Function | DNA binding, DNA-directed DNA polymerase activity, |

| | |
|---|---|
| INTERPRO | DNA polymerase, family X, beta-like, DNA-directed DNA polymerase X, Helix-hairpin-helix DNA-binding motif, DNA polymerase lambda, fingers domain, DNA polymerase family X, DNA polymerase family X lyase domain, |
| KEGG_PATHWAY | Base excision repair, |
| SMART | HhH1, POLXc, |
| **TriadG51590** | |
| COG_ONTOLOGY | DNA replication, recombination, and repair, |
| INTERPRO | Cryptochrome/DNA photolyase, class 1, DNA photolyase, FAD-binding/Cryptochrome, C-terminal, DNA photolyase, Rossmann-like alpha/beta/alpha sandwich fold, Cryptochrome/DNA photolyase, class 1 conserved site, |
| **TriadG51591** | |
| COG_ONTOLOGY | DNA replication, recombination, and repair, |
| INTERPRO | Cryptochrome/DNA photolyase, class 1, DNA photolyase, FAD-binding/Cryptochrome, C-terminal, DNA photolyase, N-terminal, Rossmann-like alpha/beta/alpha sandwich fold, Cryptochrome/DNA photolyase, class 1 conserved site |
| **TriadG51797** | |
| INTERPRO | Low-density lipoprotein (LDL) receptor class A repeat, Low-density lipoprotein (LDL) receptor class A, conserved site, |
| SMART | LDLa, |
| **TriadG51843** | |
| INTERPRO | DNA alkylation repair enzyme, Armadillo-type fold, |
| **TriadG51870** | |
| GOTERM Molecular Function | zinc ion binding, |
| INTERPRO | WD40 repeat, Zinc finger, RING-type, Zinc finger, RING/FYVE/PHD-type, WD40/YVTN repeat-like-containing domain, WD40-repeat-containing domain, |
| SMART | RING, WD40, |

| | |
|---|---|
| **TriadG51932** | Uknown |
| **TriadG52074** | |
| GOTERM Biological Process | microtubule-based movement, |
| GOTERM Cellular Component | kinesin complex, microtubule, |
| GOTERM Molecular Function | microtubule motor activity, ATP binding, ATPase activity, |
| INTERPRO | Kinesin, motor domain, Kinesin, motor region, conserved site, P-loop containing nucleoside triphosphate hydrolase, |
| SMART | KISc, |
| **TriadG52125** | |
| INTERPRO | Sterile alpha motif domain, Sterile alpha motif/pointed domain, |
| **TriadG52445** | |
| GOTERM Cellular Component | synaptic vesicle, integral component of membrane, |
| GOTERM Molecular Function | transporter activity, |
| INTERPRO | Synaptophysin/synaptoporin, Marvel, |
| **TriadG52757** | |
| GOTERM Molecular Function | nucleotide binding, nucleic acid binding, |
| INTERPRO | RNA recognition motif domain, Nucleotide-binding, alpha-beta plait, |
| SMART | RRM, |
| **TriadG53185** | |
| INTERPRO | Epidermal growth factor-like domain, EGF-like, laminin, EGF-like, conserved site, |
| SMART | EGF_Lam, EGF, |
| **TriadG53288** | |
| GOTERM Cellular Component | integral component of membrane, |
| INTERPRO | Protein of unknown function DUF1761, |
| **TriadG53566** | |
| INTERPRO | High mobility group (HMG) box domain, |

| | |
|---|---|
| SMART | HMG, |
| **TriadG53902** | |
| GOTERM Biological Process | double-strand break repair via nonhomologous end joining, protection from non-homologous end joining at telomere, interstrand cross-link repair, |
| GOTERM Cellular Component | nuclear chromosome, telomeric region, |
| GOTERM Molecular Function | damaged DNA binding, 5'-3' exodeoxyribonuclease activity, |
| INTERPRO | Beta-lactamase-like, DNA repair metallo-beta-lactamase, |
| KEGG_PATHWAY | Non-homologous end-joining, |
| **TriadG54493** | |
| GOTERM Molecular Function | phosphatidylinositol phosphate kinase activity, |
| INTERPRO | Phosphatidylinositol-4-phosphate 5-kinase, core, Phosphatidylinositol-4-phosphate 5-kinase, C-terminal, Phosphatidylinositol-4-phosphate 5-kinase, N-terminal domain, |
| KEGG_PATHWAY | Inositol phosphate metabolism, Phosphatidylinositol signaling system, Phagosome, |
| SMART | PIPKc, |
| **TriadG55476** | |
| GOTERM Molecular Function | nucleotide binding, nucleic acid binding, zinc ion binding, |
| INTERPRO | G-patch domain, RNA recognition motif domain, Zinc finger, RanBP2-type, Zinc finger, C2H2, Nucleotide-binding, alpha-beta plait, |
| SMART | RRM, G_patch, |
| **TriadG55661** | |
| GOTERM Biological Process | mitochondrial genome maintenance, recombinational repair, DNA strand renaturation, interstrand cross-link repair, |
| GOTERM Cellular Component | mitochondrial chromosome, mitochondrial nucleoid, |
| GOTERM Molecular Function | DNA binding, |
| INTERPRO | Mitochondrial genome maintenance MGM101, |
| **TriadG55798** | |

| | |
|---|---|
| COG_ONTOLOGY | Translation, ribosomal structure and biogenesis, |
| GOTERM Biological Process | exonucleolytic trimming to generate mature 3'-end of 5.8S rRNA from tricistronic rRNA transcript (SSU-rRNA, 5.8S rRNA, LSU-rRNA), nuclear-transcribed mRNA catabolic process, exonucleolytic, 3'-5', U1 snRNA 3'-end processing, U4 snRNA 3'-end processing, U5 snRNA 3'-end processing, exonucleolytic nuclear-transcribed mRNA catabolic process involved in deadenylation-dependent decay, nuclear mRNA surveillance, nuclear polyadenylation-dependent rRNA catabolic process, nuclear polyadenylation-dependent tRNA catabolic process, nuclear polyadenylation-dependent mRNA catabolic process, |
| GOTERM Cellular Component | nuclear exosome (RNase complex), cytoplasmic exosome (RNase complex), |
| GOTERM Molecular Function | AU-rich element binding, |
| INTERPRO | Exoribonuclease, phosphorolytic domain 1, Exoribonuclease, phosphorolytic domain 2, Ribosomal protein S5 domain 2-type fold, PNPase/RNase PH domain, |
| KEGG_PATHWAY | RNA degradation, |
| **TriadG56020** | |
| GOTERM Cellular Component | integral component of membrane, |
| **TriadG56088** | Uknown |
| **TriadG56259** | |
| GOTERM Biological Process | regulation of actin filament polymerization, |
| INTERPRO | FCH domain, Src homology-3 domain, |
| SMART | FCH, SH3, |
| **TriadG56514** | |
| GOTERM Biological Process | protein ubiquitination involved in ubiquitin-dependent protein catabolic process, |
| GOTERM Cellular Component | nucleus, cytoplasm, |
| GOTERM Molecular Function | ubiquitin-protein transferase activity, ligase activity, |
| INTERPRO | HECT, |

| | |
|---|---|
| SMART | HECTc, |
| **TriadG56741** | |
| INTERPRO | Hpc2-related domain, Ubinuclein middle domain, |
| **TriadG56959** | |
| INTERPRO | Armadillo-type fold, |
| **TriadG57189** | |
| GOTERM Biological Process | actin filament organization, |
| GOTERM Cellular Component | actin cytoskeleton, |
| **TriadG57566** | |
| GOTERM Biological Process | DNA replication, DNA recombination, DNA ligation involved in DNA repair, DNA biosynthetic process, |
| GOTERM Cellular Component | nucleus, mitochondrion, |
| GOTERM Molecular Function | DNA binding, DNA ligase (ATP) activity, ATP binding, zinc ion binding, |
| INTERPRO | DNA ligase, ATP-dependent, Zinc finger, PARP-type, DNA ligase, ATP-dependent, DNA ligase, ATP-dependent, Nucleic acid-binding, OB-fold, Zinc finger, C2H2, APLF-like, |
| KEGG_PATHWAY | Base excision repair, |
| SMART | SM01336, |
| **TriadG57629** | |
| GOTERM Biological Process | protein import into nucleus, |
| GOTERM Cellular Component | nucleus, cytoplasm, integral component of membrane, |
| **TriadG58144** | |
| GOTERM Cellular Component | integral component of membrane, |
| GOTERM Molecular Function | sodium channel activity, |
| INTERPRO | Na+ channel, amiloride-sensitive, |
| **TriadG58306** | |
| GOTERM Biological Process | acyl-CoA metabolic process, palmitic acid biosynthetic process, |

| | |
|---|---|
| GOTERM Cellular Component | cytosol, |
| GOTERM Molecular Function | palmitoyl-CoA hydrolase activity, |
| INTERPRO | Thioesterase superfamily, |
| KEGG_PATHWAY | Fatty acid elongation, Biosynthesis of unsaturated fatty acids, |
| **TriadG58689** | |
| GOTERM Biological Process | cell surface receptor signaling pathway, G-protein coupled receptor signaling pathway, |
| GOTERM Cellular Component | integral component of plasma membrane, |
| GOTERM Molecular Function | G-protein coupled receptor activity, |
| INTERPRO | G protein-coupled receptor, rhodopsin-like, GPCR, rhodopsin-like, 7TM, |
| **TriadG59637** | |
| GOTERM Biological Process | cell surface receptor signaling pathway, G-protein coupled receptor signaling pathway, |
| GOTERM Cellular Component | integral component of plasma membrane, |
| GOTERM Molecular Function | G-protein coupled receptor activity, |
| INTERPRO | G protein-coupled receptor, rhodopsin-like, GPCR, rhodopsin-like, 7TM, |
| **TriadG60167** | |
| GOTERM Cellular Component | mitochondrion, |
| INTERPRO | Protein of unknown function DUF1640, |
| **TriadG60371** | |
| GOTERM Biological Process | peptidyl-serine phosphorylation, intracellular signal transduction, protein autophosphorylation, |
| GOTERM Cellular Component | nucleus, cytoplasm, |
| GOTERM Molecular Function | calmodulin-dependent protein kinase activity, calmodulin binding, ATP binding, calcium-dependent protein serine/threonine kinase activity, |
| INTERPRO | Protein kinase, catalytic domain, Protein kinase-like domain, |
| **TriadG60751** | Uknown |
| **TriadG60882** | |

| | |
|---|---|
| INTERPRO | Tetratricopeptide-like helical, Tetratricopeptide repeat-containing domain, Tetratricopeptide repeat, |
| SMART | TPR, |
| **TriadG61077** | |
| GOTERM Biological Process | adenosine catabolic process, hypoxanthine salvage, inosine biosynthetic process, |
| GOTERM Cellular Component | cytosol, |
| GOTERM Molecular Function | adenosine deaminase activity, |
| INTERPRO | Adenosine/AMP deaminase domain, Adenosine/adenine deaminase, |
| **TriadG61611** | |
| INTERPRO | Death-like domain, |
| **TriadG61626** | |
| GOTERM Biological Process | telomere maintenance, double-strand break repair via nonhomologous end joining, DNA recombination, DNA duplex unwinding, cellular response to gamma radiation, cellular response to X-ray, |
| GOTERM Cellular Component | nuclear chromosome, telomeric region, Ku70:Ku80 complex, |
| GOTERM Molecular Function | damaged DNA binding, ATP-dependent DNA helicase activity, telomeric DNA binding, |
| INTERPRO | von Willebrand factor, type A, SAP domain, Ku70/Ku80 C-terminal arm, Ku70/Ku80, N-terminal alpha/beta, Ku70/Ku80 beta-barrel domain, Ku70, SPOC like C-terminal domain, Ku70, bridge and pillars domain, |
| KEGG_PATHWAY | Non-homologous end-joining, |
| PIR_SUPERFAMILY | Ku DNA-binding complex, Ku70 subunit [Parent=PIRSF800001], |
| SMART | SAP, Ku78, |
| **TriadG62277** | Uknown |
| **TriadG62514** | Uknown |
| **TriadG62635** | |
| GOTERM Cellular Component | synaptic vesicle, integral component of membrane, |

| | |
|---|---|
| GOTERM Molecular Function | transporter activity, |
| INTERPRO | Synaptophysin/synaptoporin, Marvel, |
| **TriadG62773** | |
| GOTERM Biological Process | Arp2/3 complex-mediated actin nucleation, |
| GOTERM Cellular Component | Arp2/3 protein complex, |
| GOTERM Molecular Function | ATP binding, |
| INTERPRO | Actin-related protein, Actin/actin-like conserved site, Actin-related protein 2 (Arp2), |
| **TriadG63052** | |
| **TriadG63511** | |
| GOTERM Biological Process | methylglyoxal catabolic process to D-lactate via S-lactoyl-glutathione, |
| GOTERM Molecular Function | glyoxalase III activity, |
| **TriadG6927** | |
| GOTERM Cellular Component | integral component of membrane, |
| INTERPRO | Vitamin K epoxide reductase, |
| KEGG_PATHWAY | Ubiquinone and other terpenoid-quinone biosynthesis, |
| SMART | VKc, |
| **TriadG8412** | Uknown |
| **TriadG9891** | |
| INTERPRO | Heat shock protein 70 family, |

**Table 2S.** Coverage of whole genome sequencing. There is a substantial difference between the coverage of the X-ray exposed parental and extruded samples. This is mostly caused by the limited-dying number of cells available in the extrusions. We found an average of 1847.8 total mutations per Mb (group 1 and 2) and 38.8 mutations per Mb in animals exposed to X-rays after 2 years.

| Sample | Coverage (%) | Mbp | Total mutations | Non-synonym. mutations | Total mutations/Mbp | Non-synonym. mutations/Mbp |
|---|---|---|---|---|---|---|
| Parental 1 | 10.1 | 10.8 | 41255 | 1607 | 3820 | 148.8 |
| Small asymm. fission 1 | 9.3 | 10 | 39073 | 1471 | 3907 | 147.1 |
| Extrusion 1 | 2.7 | 2.9 | 4028 | 194 | 1389 | 145.52 |
| Parental 2 | 28.7 | 30.7 | 1071 | 59 | 34.9 | 1.9 |
| Extrusion 2 | 7.6 | 8.1 | 712 | 23 | 87.9 | 2.8 |
| Untreated control animals | 89.1 | 94.1 | - | - | - | - |
| Animals exposed to X-rays after 2 years | 89 | 94 | 3636 | 253 | 38.7 | 2.7 |

**Table 3S.** Sequences of primers (5'-3') of *T. adhaerens* genes and their human orthologs (as indicated in parentheses), *TriadT64020* (*GAPDH*), *TriadG64021* (*TP53*) and *TriadG54791* (*MDM2*).

| Gene | Homology | Strand | Sequence (5' - 3') | Amplificon Size (bp) |
|---|---|---|---|---|
| ***TriadT64020*** | *GAPDH* | Forward | AAAGGGTGGCGTAGATGTTG | 93 |
| | | Reverse | TGCCATGTGTCGAATCGTAT | |
| ***TriadG64021*** | *TP53* | Forward | GCTGCAAAATGCTGTAACGA | 150 |
| | | Reverse | ACATTCGCAAGATGACCACA | |
| **TriadG54791** | *MDM2* | Forward | TGGAAGCAGAGAATTGAACG | 118 |
| | | Reverse | TTGCATCAATTCGGCATTAG | |
